# Mitochondrial coenzyme Q redox balance restrains dopaminergic neuron activity to promote early-life sleep in *Drosophila*

**DOI:** 10.64898/2026.02.22.707308

**Authors:** Jeffrey B. Rosa, Hayle H. Kim, Jianing Yang, Jenny Luong, Rishith Ramamurthy, Peyton Yee, Allen Yan, Anyara Rodriguez, Joseph A. Baur, Marni J. Falk, Eiko Nakamaru-Ogiso, Matthew S. Kayser

## Abstract

Sleep is especially abundant during early life, yet the cellular mechanisms that maintain elevated juvenile sleep drive remain poorly understood. To identify cell-intrinsic regulators of juvenile sleep, we profiled gene expression in *Drosophila* dopaminergic neurons (DANs) at different ages and performed a targeted RNAi screen of genes with enriched juvenile expression. We found that the magnitude of mitochondrial complex I (MCI) disruption produced strikingly distinct physiological outcomes. Severe MCI loss-of-function caused profound mitochondrial dysfunction, reduced dopaminergic activity, and locomotor impairment. In contrast, partial MCI inhibition left overall mitochondrial function intact but instead reduced sleep, with disproportionately strong effects on juvenile sleep fragmentation and depth, consistent with a gain of dopaminergic signaling. Multiple genetic manipulations that deplete the reduced coenzyme Q pool (CoQH₂) converged on a phenotype of increased dopaminergic activity and wakefulness, indicating that coenzyme Q redox balance itself regulates dopaminergic output. These findings support a model in which CoQH_2_-driven reverse electron transfer (RET) at MCI restrains dopaminergic arousal. By dissociating mitochondrial redox signaling from catastrophic mitochondrial failure, our work identifies a physiological role for coenzyme Q-dependent signaling in the regulation of neural activity and behavioral state. More broadly, these findings suggest that sleep phenotypes may serve as sensitive indicators of emerging mitochondrial dysfunction and reveal a potential link between the regulation of early-life sleep and later dopaminergic vulnerability.

## Introduction

Sleep characteristics change over the lifespan. Across the animal kingdom, juvenile animals exhibit longer and deeper sleep states than their adult counterparts. In humans, newborns spend up to 70% of the day asleep, and, compared to adults, newborn sleep is disproportionately enriched in rapid eye movement (REM) sleep(1, 2),. At a molecular level, human genome wide association studies in adolescents and adults point to unique genetic variants impacting sleep quality at these different ages(3). Likewise, genetic regulators of sleep appear to be at least partially distinct in early life compared to maturity in animal models(4, 5). These findings suggest that early-life sleep is under distinct genetic and neural control, resulting in its unique attributes.

The ontogenetic hypothesis of sleep postulates an essential role for juvenile sleep in brain and behavior maturation(2). Early-life sleep disruption in both invertebrates and vertebrates causes later deficits in social behavior associated with synaptic abnormalities in the brain(4–7). These observations indicate that early-life sleep could play instructive and/or permissive roles during typical brain development.

Moreover, early-life sleep disturbances are a convergent comorbidity of neurodevelopmental disorders, with the severity of sleep disturbances correlating with the severity of social deficits(8, 9) Thus sleep disruption in children with atypical neural development may be a driver of later behavioral and cognitive deficits.

The fruit fly *Drosophila melanogaster* has been a powerful model system to investigate the genetic and circuit regulation of early-life sleep. Compared to their mature adult (∼6-10 days old) counterparts, juvenile adult flies (0-1 day old) fall asleep faster, sleep more, and experience deeper sleep states(5, 10, 11). Previous work has found that high sleep drive in juvenile flies is facilitated by relatively hypoactive dopaminergic neurons, which normally promote arousal(5). During the first week of adult life, dopaminergic activity increases, leading to a reduction in sleep duration, in part through greater inhibition of sleep-promoting neurons(5, 12). Work in mammalian models likewise implicates dopamine signaling in changes to sleep across development(4). However, the cell intrinsic processes acting in DANs to restrict their activity and facilitate excess sleep in juvenile periods remain unknown.

Here, using transcriptomics-guided genetic screening in *Drosophila* dopaminergic neurons, we identify mitochondrial Complex I (MCI) as a regulator of dopaminergic arousal signaling that preferentially influences juvenile sleep. Strikingly, distinct degrees of MCI perturbation produce opposing behavioral and cellular outcomes: severe disruption causes dopaminergic hypoactivity and locomotor impairment, whereas partial perturbation leaves overall mitochondrial function intact but selectively reduces sleep by augmenting dopaminergic activity. Based on multiple orthogonal manipulations, we propose that sleep loss is caused by coenzyme Q redox imbalance --specifically, depletion of reduced coenzyme Q/ubiquinol. We propose a model in which reverse electron transfer (RET)-related mitochondrial signaling restrains dopaminergic neuron activity to promote juvenile sleep. Finally, we find that partial MCI perturbation is associated with age-dependent dopaminergic neuron loss and shortened lifespan, linking mitochondrial regulation of early-life sleep to later neuronal vulnerability.

## Results

### Transcriptomic analysis of juvenile and adult dopaminergic neurons

Maturation of sleep patterns across early life involves changes to dopaminergic neuron (DAN) activity, as high juvenile sleep drive requires dopaminergic hypoactivity(5). We hypothesized that transcriptional differences between juvenile and mature adult DANs might play a role in this functional change. We used fluorescence activated cell sorting to isolate DANs from brains of juvenile (0-1 day old) and mature (6-10 day old) adult flies (**Figure 1A**). Focusing on genes with higher expression in juvenile DANs, we observed an enrichment of GO terms for metabolic pathways, vacuolar ATPase components, redox biology and synaptic signaling (**Supplemental Figure 1A**). Notably, components of the oxidative phosphorylation pathway were the most significantly enriched using both GO analysis and KEGG pathway analysis (**Supplemental Figure 1 A,B)**, including multiple regulators of the Coenzyme Q/Ubiquinone redox state, suggesting heightened demand for metabolic gene expression. To test this possibility, we measured the NADH:CoQ oxidoreductase activity of Complex I and II in whole flies at different ages. Compared to their mature (9-10 days old) counterparts, 1 day-old juvenile wild-type (*iso^31^*) flies exhibited ∼50% higher Complex II activity and a trend toward higher Complex I activity (∼20% higher; **Supplemental Figure 1C**), consistent with altered electron transport chain function in juvenile stages. Together, these findings suggest that juvenile flies possess a distinct metabolic state, marked by increased expression and activity of electron transport chain components.

**Figure 1:**
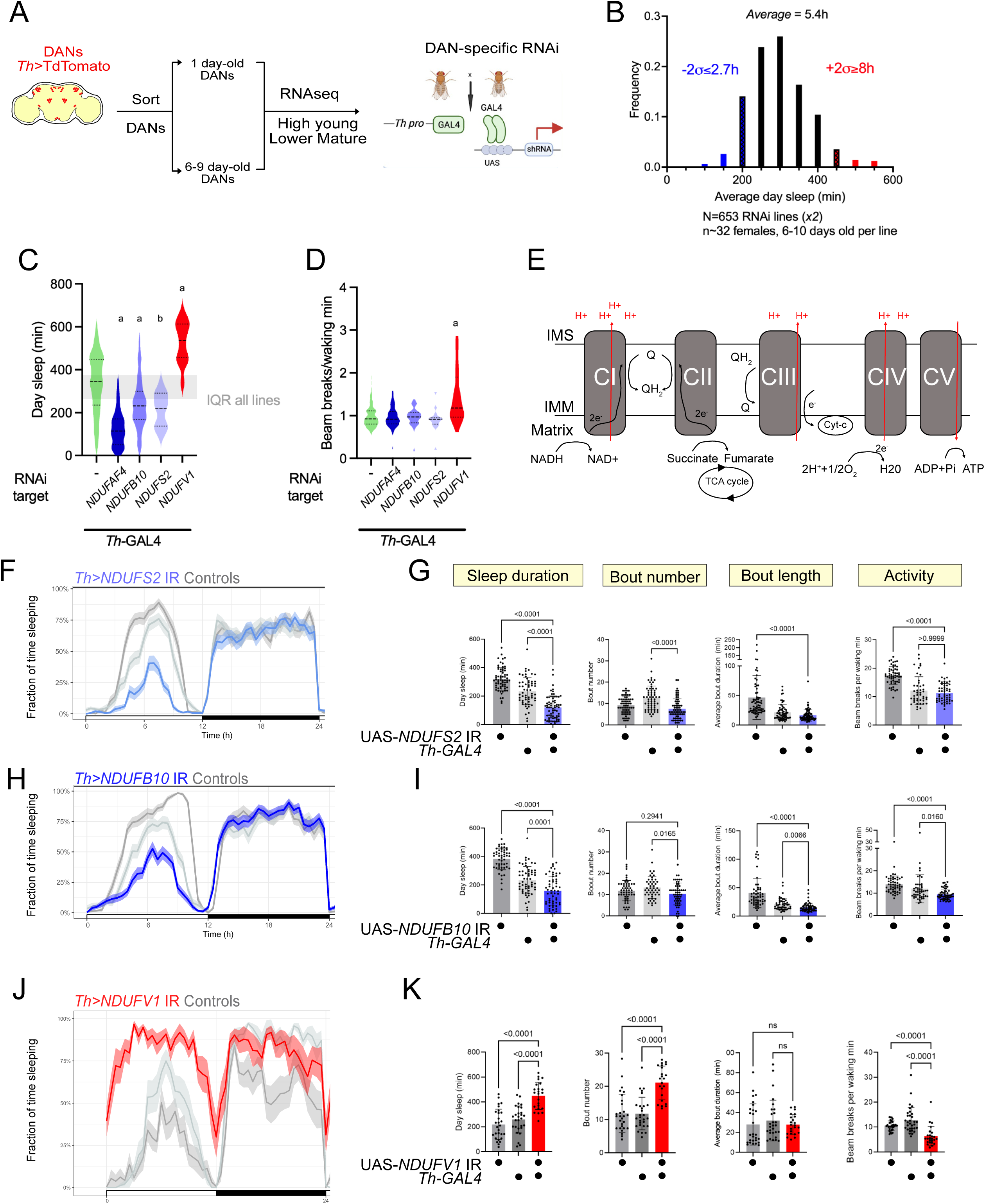
Mitochondrial Complex I loss-of-function in dopaminergic neurons causes aberrant sleep. (A) Workflow of RNAi-based sleep screen using high throughput activity monitoring. (B) Average daytime (ZT0-12) sleep duration from Th>RNAi screen conducted in 6-10 day old adult flies (n∼32 per genotype, 2 biological replicates). Th>RNAi genotypes with day sleep values ≥2 standard deviations from mean were selected as hits. Short-sleepers represented in blue, long-sleepers in red. (C) Knock-down of Mitochondrial Complex I (MCI) subunits caused either sleep loss (blue) or excess sleep (red). Grey shading represents the range of the middle 50% of RNAi lines tested. (a: p<0.0001 Kruskal-Wallis test with Dunn’s multiple testing comparisons; b: p=0.0032 Kruskal-Wallis with Dunn’s multiple testing comparisons; all comparisons to Th>+ control) (D) Waking activity of Th>MCI RNAi conditions as assayed using single-beam activity monitors. (a: p<0.0001 Kruskal-Wallis with Dunn’s multiple testing comparisons, compared to *Th*>+). (E) MCI is an entry point for reducing equivalents from glycolysis and the TCA cycle, oxidizing NADH and shuttling electrons to the mobile electron carrier Coenzyme Q, coupled to H+ translocation. (F) Sleep per 30 min windows across a 12h:12h light: dark period measured using multi-beam activity monitoring: *Th>NDUFS2* IR trace (average with 95% confidence interval shaded) is in blue and controls in grey. (G) Sleep metrics during ZT0-12 for *Th>NDUFS2* IR and controls. (H) Sleep per 30 min windows across a 12h:12h light:dark period measured using multi-beam activity monitoring: Th>*NDUFB10* IR trace is in blue and controls in grey. (I) Sleep metrics during ZT0-12 for *Th>NDUFB10* IR flies (blue) and controls (grey). (J, K) When assayed using multi-beam activity monitoring, Th>*NDUFV1* IR flies (red) exhibited increased quiescence associated with reduced locomotor activity. P values computed using either Kruskal-Wallis tests with Dunn’s multiple-testing comparisons or Welch ANOVA with Dunnett’s T3 multiple-testing comparisons.

Based on the DAN transcriptional analysis, we next sought to assess the role of DEGs in sleep. To enrich for genes with potential large effects in regulating DAN activity, we narrowed our focus to genes with both i) higher juvenile expression and ii) high expression across all conditions (above-median base expression) (**Supplemental Figure 1C**). We used a *Th-GAL4* driver to express dsRNAs/shRNAs against these genes in dopaminergic neurons, combined with high throughput activity monitoring to track sleep in 6-10 day old female flies (n∼32 flies, across two biological replicates) under 12:12h light-dark conditions **(Figure 1A).** Mature adult, and not juvenile flies, were used to increase the throughput of the screen. We focused on changes to day sleep, since ontogenetic sleep differences are most pronounced for this period(5, 10). Across all RNAi lines (n=653), flies slept on average ∼5.4h during the day **(Figure 1B).** Using a two standard deviation (2SD) cut-off, we identified 6 *Th>*RNAi genotypes sleeping 2.7h or less as short sleepers and 23 *Th >*RNAi genotypes sleeping 8h or more as long-sleepers **(Figure 1B).**

### Disruption of Mitochondrial Complex I causes abnormal sleep

Given that low DAN activity is a key feature of elevated juvenile sleep drive(4, 5), we reasoned that short-sleep lines recovered from our screen would enrich for LOF conditions causing excess DAN activity. Our shortest sleeping hit (2.1 +/- 1.4h of day sleep) was an RNAi line targeting *NDUFAF4*, an assembly factor for Mitochondrial Complex I (MCI) **(Figure 1C).** Moreover, we uncovered two additional RNAi lines targeting integral MCI subunits (*NDUFB10* and *NDUFS2*) as causing reduced sleep, with 3.9 +/- 1.9h of day sleep for *Th>NDUFB10* IR and 3.6 +/- 1.3h for *Th>NDUFS2* IR flies **(Figure 1C).** None of these RNAi lines affected waking locomotor activity **(Figure 1D)**, arguing against nonspecific motor changes. In *Drosophila,* MCI consists of 43 subunits organized into an inner membrane-associated domain and a hydrophilic arm protruding into the matrix(13). The hydrophilic domain initiates electron transport by accepting 2 electrons from reduced nicotinamide adenine dinucleotide (NADH) and shuttling them toward Coenzyme Q in the inner mitochondrial membrane. This transfer is coupled to H^+^ translocation against its electrochemical gradient **(Figure 1E).**

NDUFAF4 promotes the assembly of the large holoenzyme complex from smaller subunits(14). NDUFS2 contains an Fe-S cluster that directly participates in the MCI redox cycle and also contributes residues to the Coenzyme Q binding site. NDUFB10 encodes an accessory subunit of the membranous arm of MCI and is necessary for MCI assembly and function(15, 16). Collectively, the screen uncovered converging lines of evidence that loss of MCI function in DANs causes sleep loss.

### Partial versus strong MCI loss-of-function explains divergent sleep phenotypes

Neurons have high ATP requirements due to the elaboration and maintenance of their neurites, maintenance of their resting membrane potential, and synaptic vesicle cycling. It was therefore surprising to find MCI loss-of-function (LOF) caused *hyper*arousal, suggestive of elevated DAN activity. However, we also recovered an RNAi line targeting the NADH-binding subunit of MCI, NDUFV1, as one of our longest-sleeping flies (8.8 +/- 1.6h of day sleep, **Figure 1C**), consistent with DAN hypofunction. What accounts for these opposing sleep phenotypes?

First, to validate our screen hits, we repeated the MCI RNAi experiments using additional genetic controls and higher spatial resolution sleep monitoring. We replicated the mature adult day sleep loss seen in *Th>NDUFB10* IR and *Th>NDUFS2* IR flies **(Figure 1F-I).** Night sleep metrics were unaffected (**Supplemental Figure 2A,B**). The *Th>NDUFAF4* IR-associated reduction in daytime sleep was significantly different compared to only one genetic control, but with many flies sleeping far less than normal (**Supplemental Figure 2D-G**). In contrast, the increased daytime quiescence observed in *Th>NDUFV1* IR flies (**Figure 1J, K)** was correlated with a reduction in locomotor activity (**Supplemental Figure 2C**), indicating that increased quiescence can be explained by general hypoactivity. As with the short-sleeping lines, night sleep measures were unaffected in *Th>NDUFV1* IR flies (**Supplemental Figure 2B**).

We next tested if the short sleeping vs hypoactive *Th>*MCI dopaminergic RNAi phenotypes reflect differential MCI LOF or subunit-specific roles. We expressed UAS- *NDUFV1* IR pan-neuronally (*elav>NDUFV1* IR) and observed a prolonged wandering third instar stage and pupal lethality. By contrast, *elav>NDUFB10* IR, *elav>NDUFS2* IR, and *elav>NDUFAF4* IR flies all survived into adulthood with no visible defects (**Table 1**). This result suggests either that neurons are differentially sensitive to loss of specific MCI regulators, or that the short-sleeping RNAi lines preserve enough MCI activity to sustain flies through pupation. To rule out the former hypothesis, we generated transgenic multiplex guide RNA flies targeting coding regions of *NDUFB10* (short-sleep), *NDUFS2* (short-sleep), and *NDUFV1* (hypoactive)(*17*). We used pan-neuronal CRISPR to ablate the integral MCI subunits *NDUFB10, NDUFS2*, and *NDUFV1* in neurons, reasoning that the Cas9-induced genetic lesions should cause a stronger LOF than RNAi. Indeed, using a pan-neuronally expressed Cas9 (*elav>Cas9),* we found that somatic neuronal mutagenesis of all three loci was adult lethal: *elav>Cas9; g-NDUFB10* and *elav>Cas9; g-NDUFS2* progeny died as pupae (**Table 1**). Thus, adult neurons do indeed require *NDUFB10* and *NDUFS2* function to complete pupation, arguing against subunit-specific phenotypes. We then used quantitative PCR to measure mRNA knock-down efficiency for each RNAi line. In line with previously published findings in muscle using identical RNAi lines(18), we found the short-sleeping MCI RNAi lines depleted their target mRNAs but to a lesser extent than the hypoactive UAS-*NDUFV1* RNAi line (**Table 1**). (**Table 1**). These results are consistent with the short-sleeping RNAi lines causing partial MCI LOF.

**Table 1.**
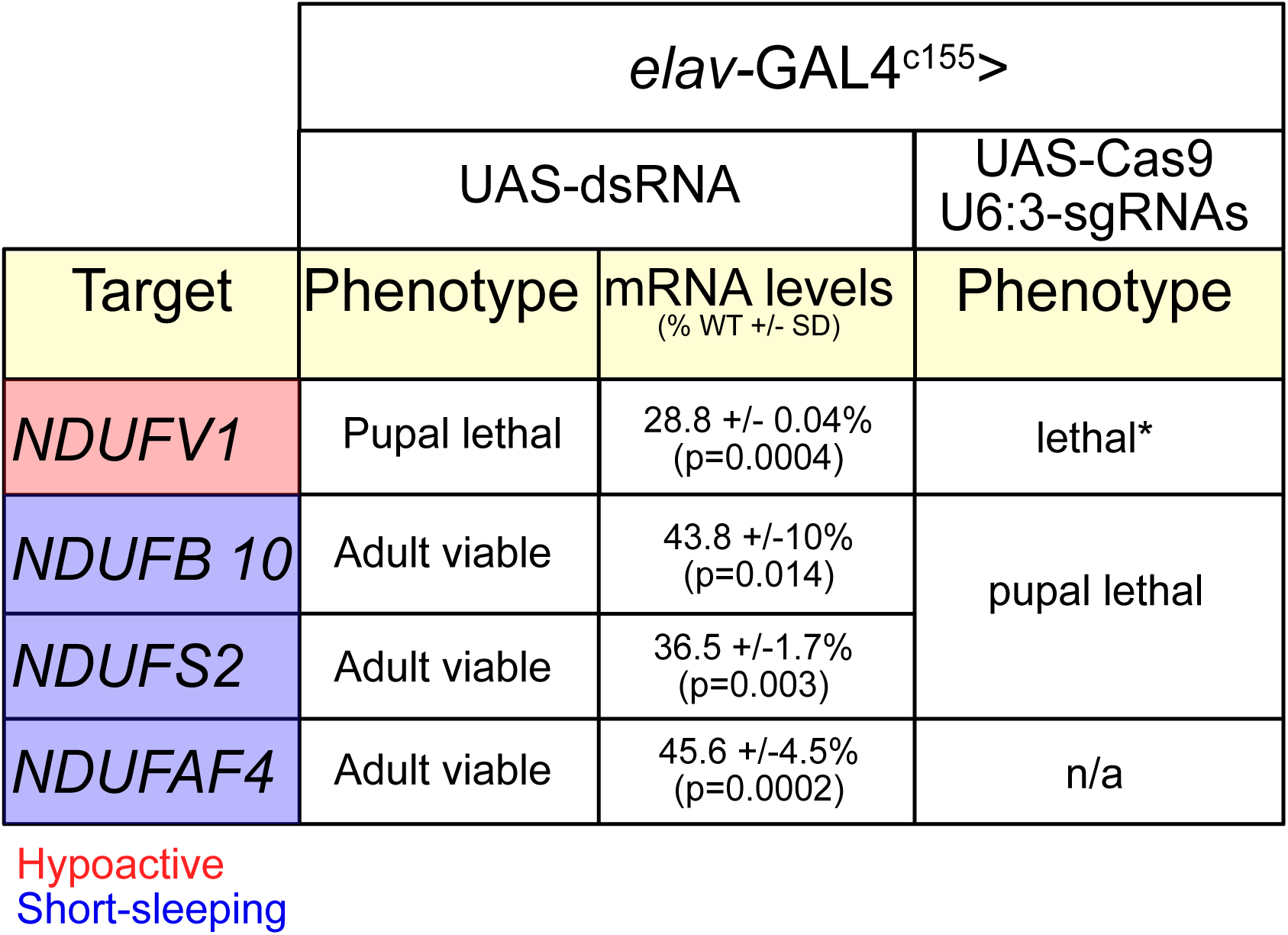
Summary of phenotypes from pan-neuronal MCI manipulations. The *elav-GAL4* pan-neuronal GAL4 driver was used to manipulate MCI subunit function either using RNAi or somatic CRISPR-mediated mutagenesis using a UAS-Cas9 transgene. *Act5C-GAL4* was used for ubiquitous expression. P values computed using Welch’s t tests. *Precise stage of lethality not determined for *elav>Cas9; g-NDUFV1*.

To address RNAi efficacy and impact differences in dopamine neurons more directly, we examined DAN morphology in *Th>NDUFB10* IR (partial LOF) and *Th>NDUFV1* IR (strong LOF) brains, focusing on the PPL1 sub cluster (**Figure 2A**) given their relevance to sleep/wake behavior(19, 20). Th>*NDUFV1* IR DAN cell bodies contained prominent cytoplasm-excluding structures not found in control or *Th>NDUFB10* IR DAN cell bodies (**Figure 2B**). These structures corresponded to swollen mitochondria not observed in DANs from other conditions (**Figure 2C,D**), indicating that mitochondrial function is more severely compromised in DANs of hypoactive *Th>NDUFV1* IR flies compared with short-sleeping *Th>NDUFB10* IR flies. Thus, while partial MCI disruption results in reduced sleep, general hypoactivity emerges as a consequence of severe mitochondrial dysfunction.

**Figure 2:**
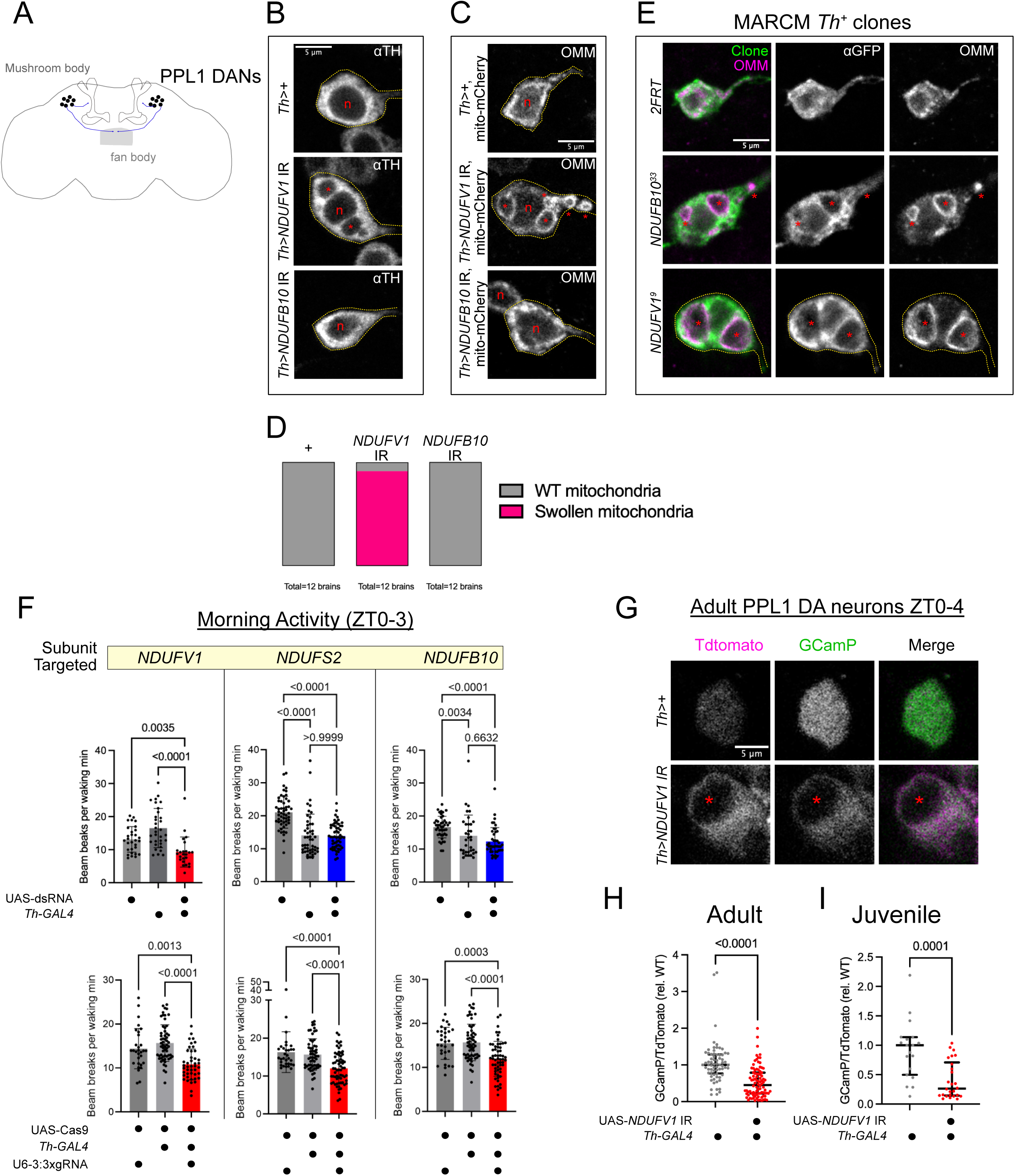
Strong MCI loss-of-function caused dopaminergic hypofunction. (A) Paired posterior lateral 1 (PPL1) dopaminergic neurons extend axons to sleep-regulatory regions in the fan body of the central complex and mushroom bodies. (B) Confocal micrographs of the cell bodies of PPL1 DANs labeled by anti-Tyrosine Hydroxylase in control (Th>+), *Th>NDUFV1* IR and *Th>NDUFB10* IR brains. DANs in *Th>NDUFV1* IR brains accumulated cytoplasm-excluding structures (asterisks) around the nucleus (n) not found in control or *Th>NDUFB10* IR brains. (C) PPL1 DANs co-expressing MCI RNAi transgenes and a mito.OMM-mCherry reporter labeling the outer mitochondrial membrane (OMM). In control and *Th>NDUFB10* IR cells, mitochondria in the cell bodies are found in a diffuse reticular network filling the cell body. In *Th>NDUFV1* IR cells, mito.OMM-mCherry labels swollen mitochondria in which the mitochondrial lumens (*) were resolvable. (D) Quantification of the penetrance of swollen mitochondria in PPL1 DANs from control (Th>+), *Th>NDUFV1* IR, and *Th>NDUFB10* IR brains. (E) CD8::GFP-labeled wild-type (*2FRT), NDUFB10^33^, and NDUFV1^9^* MARCM PPL1 clones expressing mito.OMM-mCherry. MCI null clones for either MCI subunit exhibited swollen mitochondria (asterisks), demonstrating that both NDUFB10 and NDUFV1 subunits are essential for normal dopaminergic mitochondrial morphology. (F) Waking locomotor activity during ZT0-3 for MCI RNAi (top) and MCI somatic CRISPR (bottom) manipulations in DANs. Depressed locomotor activity was observed for only strong MCI loss-of-function manipulations such as *Th>NDUFV1* IR and somatic knock-out of MCI subunits. P values computed using either Kruskal-Wallis tests with Dunn’s multiple-testing comparisons, or Welch ANOVA with Dunnett’s T3 multiple-testing comparisons. (G-I) *ex vivo* imaging of *Th-LexA>TdTomato-2A-GCamP6s; Th-GAL4> NDUFV1* IR adult (H) and juvenile (I) brains. GCamP signal is in green and TdTomato is in magenta. Strong MCI RNAi caused a significant reduction in the GCamP/TdTomato ratio, indicating reduced neuronal activity. Asterisk marks swollen mitochondria. Quantifications from n=6 brains per condition from 3 independent experiments. Data are normalized to the median value of the control brains imaged in parallel. P values computed using Mann-Whitney tests.

### *NDUFB10* is required for ongoing MCI activity and dopaminergic mitochondrial function

Mitochondrial complex I is ubiquitously expressed and supports the ATP demands of all cells in *Drosophila,* complicating functional analysis specifically in adult neurons beyond transgenic RNA interference. To clarify the function of complex I in dopaminergic mitochondrial function, we used our transgenic guide RNA flies to generate *NDUFB10* and *NDUFV1* germline mutant alleles on an FRT-bearing chromosome (**Supplemental Figure 3A, B**), allowing for the analysis of genetic mosaics. After recovering non-complementing alleles, we leveraged the essential role for oxidative phosphorylation in normal eye development to prioritize alleles for characterization(21, 22). In somatic eye clones generated using the *EGUF/Hid* approach(23), *NDUFB10^33^* caused small, glassy eyes similar to clones homozygous for a *NDUFV1* null allele (*NDUFV1^9^*; **Supplemental Figure 3C**). The *NDUFV1^9^* allele is larval lethal and results from a premature stop codon after G66, truncating the open reading frame upstream of the flavin adenine mononucleotide (FMN)-binding domain and Fe-S clusters that mediate the catalytic activity of NDUFV1 (**Supplemental Figure 3A**). Since NDUFV1 initiates electron transfer at MCI from NADH, *NDUFV1^9^* is likely to abolish most MCI activity. *NDUFB10^33^* is also larval-lethal and carries a 2-nucleotide deletion leading to a missense polypeptide N-terminal in a conserved disulfide bridge and lacking the endogenous stop codon (**Supplemental Figure 3B, E**). Mutations abolishing this disulfide bridge in humans block NDUFB10/NDUFB10 protein import into mitochondria and cause severe neonatal MCI deficiency(24).

Using this approach, we serendipitously recovered a temperature sensitive allele, named *NDUFB10^ts^*. This allele is an in-frame deletion of 4 amino acids (Y93-P96) in the linker region between the two disulfide-bonded alpha helices (**Supplemental Figure 3B,E**). *NDUFB10^ts^ e*ye clones appeared phenotypically normal at 25°C but were severely disrupted at 29°C (**Supplemental Figure 3C**). Moreover, *NDUFB10^ts^* failed to complement *NDUFB10^33^* larval lethality at 25°C or higher but could fully rescue *NDUFB10^33^* lethality at 22°C (**Supplemental Figure 3D**). Based on a high-confidence AlphaFold structure(25) of NDUFB10, Y93-P96 mediate several critical elements of tertiary structure of the disulfide-bonded alpha helices (α2 and α3): E94 in the α2-α3 linker is hydrogen-bonded to K101 in α3; the P96 side chain generates the bend of the linker region and the carbonyl group of its backbone is H-bonded with the conserved H98 side chain in α3 (**Supplemental Figure 3E**). Lower rearing temperatures could allow the stabilization of the α2 and α3 helices in the absence of these non-covalent interactions.

With these genetic tools, we were able to bypass early requirements for essential MCI subunits and directly test functional requirements in adults. First, to determine how *NDUFB10* gene dosage impacts Complex I activity, we purified whole-animal mitochondria from *NDUFB10^+/+^, NDUFB10^33/+^,* and *NDUFB10^33/ts^* genotypes to measure Complex I and Complex II enzymatic activity *in vitro*. All genotypes were reared at 22°C until eclosion and subsequently shifted to 25°C for ∼5 days to inactivate the *NDUFB10^ts^* allele. Flies with only one functional copy of *NDUFB10* showed no diminution in MCI activity compared to the wild-type, but the *NDUFB10^33/ts^* mutants exhibited an ∼80% decrease in Complex I activity (**Supplemental Figure 3F**), a reduction in activity that is comparable to strong loss-of-function of core catalytic subunits in other animals (26).

Electron transfer through Complex II was modestly increased in the *NDUFB10^ts/33^* mutants, likely reflecting a compensatory increase in Succinate Dehydrogenase activity to sustain CoQ reduction. We next generated GFP-labeled *NDUFB10^33^* MARCM clones of PPL1 DA neurons to examine how such a strong reduction in MCI activity affects mitochondrial morphology in DANs(27). To examine adult phenotypes, clones were induced in second instar larvae (48-66h after egg-laying), as this was the latest period we were able to recover *Th+* clones in adult central brains with appreciable frequency. We observed *Th+* clones in 2-4 day old adults with no apparent cell loss or gross branching defects (**Supplemental Figure 3G-I)**. However, in contrast to wild-type MARCM clones or *Th>NDUFB10* IR cells, *NDUFB10^33^* DANs exhibited swollen mitochondria closely resembling those seen in *NDUFV1^9^* null clones and *Th>NDUFV1* IR cells (**Figure 2E**). Combined with the biochemical data, these results indicate that mitochondrial swelling is accompanied only by severe reductions to MCI enzymatic activity. Therefore, while *NDUFB10* is essential for mitochondrial function in DANs during development and/or early adult stages, Complex I in *Th>NDUFB10* IR flies retains enough activity to preserve mitochondrial morphology, thereby uncovering an unexpected sleep phenotype.

### Genetic dissociation of locomotor and sleep phenotypes caused by MCI- deficiency in DA neurons

In flies, locomotor activity peaks at dawn and dusk (**Supplemental Figure 4A-B**, grey lines). This pattern is associated with elevated DA neuronal activity(28); specifically, the peak in morning activity depends on increased post-synaptic DA signal receptivity (29). When a TdTomato-2A-GCaMP6s reporter was expressed in wild-type DANs and imaged *ex vivo*, we observed higher steady-state GCaMP/TdTomato ratios in PPL1 DANs during ZT0-3 than ZT4-8 (**Supplemental Figure 4C**), establishing our ability to measure physiologically relevant changes in DA neuron activity. *Th>NDUFV1* IR flies exhibited a pronounced depression in the amplitude of morning and evening activity (**Figure 2F; Supplemental Figure 4A, G, H**). In *ex vivo* preparations of adult brains at ZT0-4, *NDUFV1* knock-down caused a reduction in the steady-state GCaMP/TdTomato ratio compared to control cells imaged in parallel (**Figure 2G, H**). A reduction in cytosolic Ca^2+^ levels was apparent even in juvenile (1 day old) *Th>NDUFV1* IR; *Th- LexA> TdTomato-2A-GCamP6s* brain preparations (**Figure 2I**), indicating *NDUFV1* depletion causes DAN hypofunction at or before eclosion. These findings suggest that reduced neuronal activity in *NDUFV1*-depleted DANs causes the hypoactivity phenotype.

In contrast to *NDUFV1* knockdown, short-sleeping *Th>NDUFB10* IR and *Th>NDUFS2* IR flies did not show the same pronounced suppression of morning and evening locomotor peaks (**Figure 2F** and **Supplemental Figure 4B,G,H**). Thus, partial MCI perturbation does not recapitulate the locomotor hypoactivity caused by strong MCI loss-of-function. To rule out the possibility that the NDUFB10 and NDUFS2 subunits were simply not required in DANs for normal locomotion, we used Th-specific Cas9 to genetically ablate these loci and assessed resultant locomotor phenotypes. Cas9 mutagenic activity was verified in our neurons of interest using a validated guide RNA targeting the GFP coding sequence(30) (**Supplemental Figure 4D-F**). We found that genetic ablation of MCI in DANs produced morning and evening hypoactivity regardless of the subunit targeted (**Figure 2F** and **Supplemental Figure 4G, H**). These data suggest that broad dopamine-dependent locomotor dysfunction emerges primarily under conditions of severe MCI impairment.

### Partial MCI inhibition augments wake-promoting DA signaling primarily during the day

The day-specific sleep loss observed in *Th>NDUFB10* IR and *Th>NDUFS2* IR flies suggests that partial MCI inhibition might interact with environmental light cues or time-of-day-dependent arousal state. Light dampens the wake-promoting effect of DA(31). Consistent with this idea, when flies were transferred to constant darkness (DD), control animals exhibited a marked reduction in subjective daytime sleep (CT0-12) relative to the preceding light-dark cycle (ZT0-12)(**Figure 3A,B**). This sleep loss was driven primarily by an increased probability of transitioning out of sleep [P(wake)] with comparatively little change in the probability of transitioning from wake to sleep [P(doze)] (**Figure 3D,E; Supplemental Figure 5C,D**), indicative of enhanced arousal and shallower sleep. Because DA promotes wakefulness principally through increasing P(wake)(27), these findings are consistent with disinhibition of a daytime DA-mediated arousal signal upon removal of the light cue. Sleep loss during CT12-24 was comparatively modest and variable, suggesting this wake-promoting signal is strongest during the subjective day (**Supplemental Figure 5E,F**).

**Figure 3:**
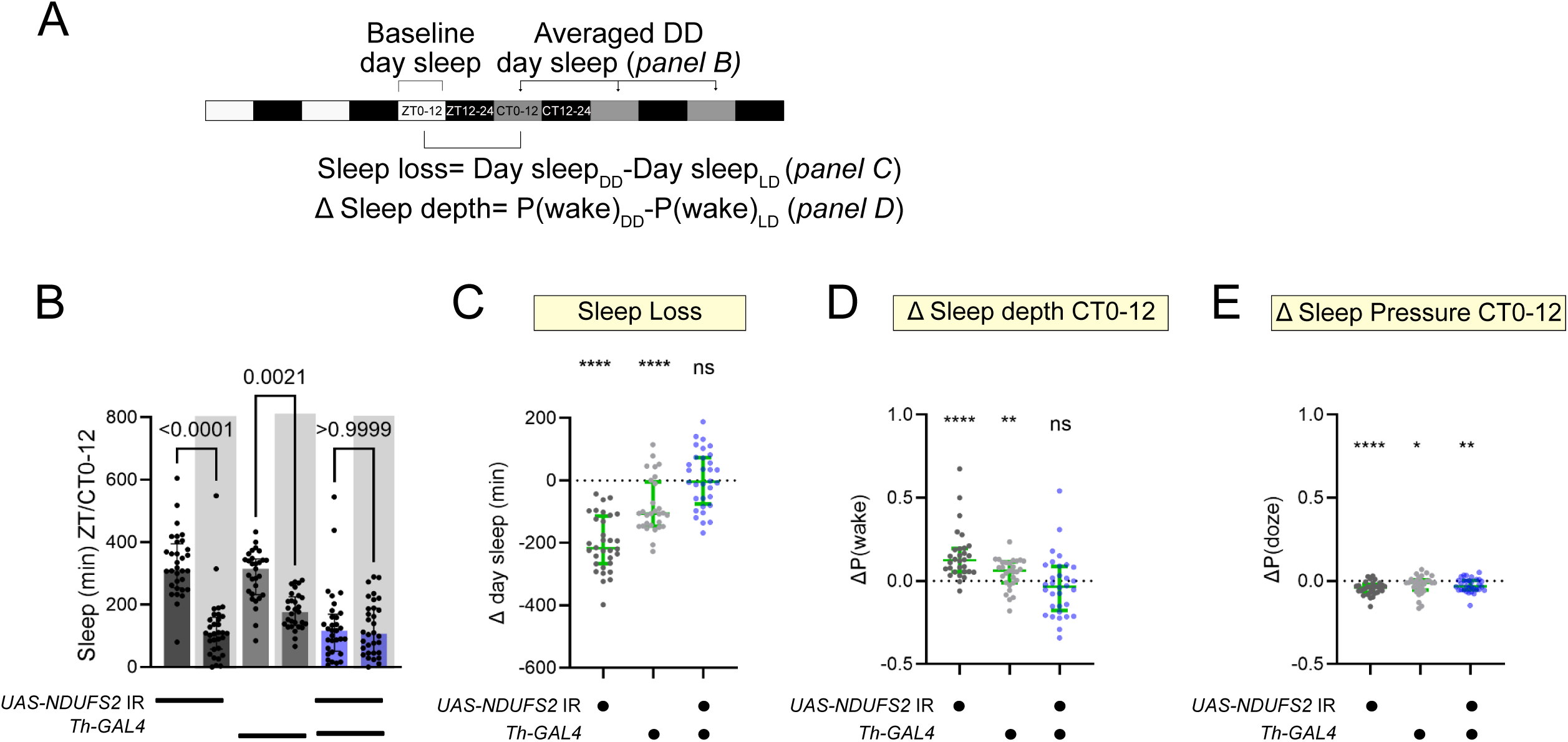
Disinhibition of DA signaling by withdrawal of the light cue modifies wild-type sleep, but not MCI-deficient, flies. (A) Light schedule for assaying sleep phenotypes in constant darkness. (B) Comparison of baseline day sleep with the light cue to subjective day sleep without the lights on (shaded, averaged across 3 successive days). *Th>NDUFS2* IR flies still exhibited short sleep under constant darkness. Control flies exhibited significant sleep loss after withdrawal of the light cue, which was not observed in *Th>NDUFS2* IR flies. (C) Change in day sleep [Day sleep(DD)-Day sleep (LD)] plotted for each fly in the experiment in B. Values < 0 (dotted line) indicate loss of sleep. *NDUFS2* knock-down abrogated sleep loss due to withdrawal of the light cue. (D) Change in P(wake) [P(wake)(DD)-P(wake)(LD)] plotted for each fly in panel B. Sleep loss upon withdrawal of the light cue was driven by an increase in P(wake)--an effect attenuated *NDUFS2* knock-down. (E) Change in P(doze) [P(doze)(DD)-P(doze)(LD)] plotted for each fly in panel B. Withdrawal of the light cue caused a slight but significant decrease in P(doze) that was not affected by *NDUFS2* knock-down. P values calculated by Wilcoxon signed rank tests (* P<0.05, ** P<0.01, **** p<0.0001 compared to Δ=0).

Interestingly, sleep in *Th>NDUFS2 IR* and *Th>NDUFB10 IR* flies during the daytime light period (ZT0-12) closely resembled sleep in control flies during the same interval under DD conditions (CT0-12) (**Figure 3B; Supplemental Figure 5A**).

Moreover, unlike controls, partial MCI knockdown flies failed to exhibit further daytime sleep loss when transferred to DD. Individual knockdown flies also lost the stereotyped sleep response to removal of the light cue and instead exhibited highly variable responses (**Figure 3C,D; Supplemental Figure 5B,C**). Together, these findings argue against a simple floor effect and instead suggest that partial MCI inhibition renders flies relatively insensitive to the wake-promoting state unmasked by withdrawal of the light cue. We therefore propose that partial MCI inhibition in DANs mimics the effect of removing the light cue by augmenting daytime DA-mediated arousal signaling, providing a likely explanation for the stronger daytime sleep phenotypes observed across our MCI perturbations.

### Partial MCI inhibition reduces juvenile sleep depth

DANs are normally less active in juvenile flies(5), facilitating increased sleep during early life. We therefore asked whether partial MCI inhibition in juvenile DANs disrupts characteristic features of juvenile sleep. Under standard LD conditions, 1 day-old Th>NDUFS2 IR, Th>NDUFB10 IR, and Th>NDUFAF4 IR flies all exhibited daytime sleep loss (**Figure 4A,E; Supplemental Figure 6A**), while nighttime effects were modest or absent. Although partial MCI inhibition also reduced sleep in mature adults, juvenile flies exhibited prominent fragmentation phenotypes not consistently observed later in life. Specifically, all three MCI perturbations shortened sleep bout duration during both day and night (**Figure 4C,D,G,H; Supplemental Figure 6C-G**), indicating a disruption of the deeper, more consolidated sleep characteristic of juvenile animals.

**Figure 4:**
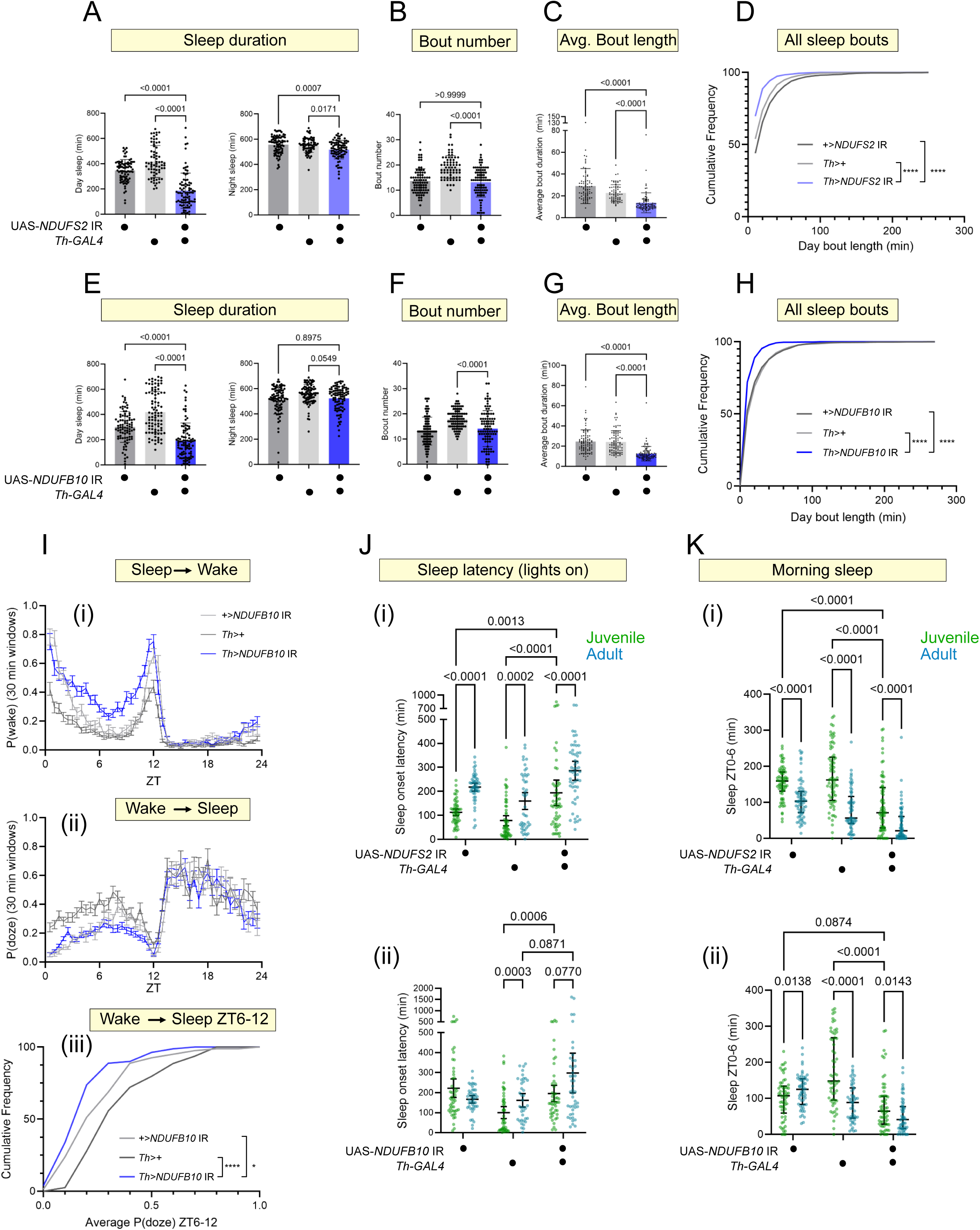
Partial MCI loss-of-function impaired juvenile sleep depth leading to fragmented sleep. (A) Day and night sleep duration in juvenile *Th>NDUFS2* IR flies (blue) and controls (grey). (B-D) Day sleep metric in juvenile *Th>NDUFS2* IR flies and controls. Sleep loss during the day was caused by reduced sleep bout duration (C,D, **** P<0.0001). (E) Day and night sleep duration in juvenile *Th>NDUFB10* IR flies (blue) and controls (grey). (F-H) Daytime sleep metric of juvenile *Th>NDUFB10* IR flies and controls. Sleep loss was caused by reduced sleep bout duration (G,H, **** p<0.0001). P values computed using either Kruskal-Wallis tests with Dunn’s multiple-testing comparisons or Welch ANOVA tests with Dunnett’s T3 multiple-testing comparisons. (I) P(wake) (i) and P(doze) (ii) calculated across 30 min intervals (mean +/- SEM) across ZT0-24 in *Th>NDUFB10* IR juvenile flies (blue) and controls (grey). Knock-down flies showed consistently increased in P(wake) during ZT0-12 compared to control flies, indicating impaired sleep depth. P(doze) was only significantly reduced in *Th>NDUFB10* IR juvenile flies during ZT6-12 (iii). P values computed using Kruskal-Wallis tests with Dunn’s multiple-testing comparisons. (J) Time to the first sleep bout after ZT0 (sleep latency) for juvenile (green) and mature (blue) *Th>NDUFS2* IR flies and genetic controls (i) and *Th>NDUFB10* IR flies and genetic controls (ii). Robust MCI activity was required to maintain relatively short sleep latencies observed in juvenile flies. (K) Sleep ontogeny phenotypes in *Th>NDUFS2* IR (i) and *Th>NDUFB10* IR (ii) flies as measured by sleep duration during ZT0-6. Juvenile flies (green dots) across all genotypes exhibited greater sleep during the first part of the day compared to their mature counterparts (blue dots), indicating sleep maturation is not impaired by partial MCI LOF. P values in J and K computed using Two-way ANOVA with mix-effects model and Tukey’s multiple comparisons tests

Juvenile flies normally exhibit reduced sleep-to-wake transitions and rapidly initiate sleep following light to dark transitions(5). Consistent with impaired juvenile sleep depth, all three partial MCI perturbations increased daytime P(wake) during ZT0- 12 (**Figure 4I(i); Supplemental Figure 6H,I**). By contrast, effects on P(doze) were less consistent across genotypes and time windows (**Figure 4I(ii-iii)**). Juvenile *Th>NDUFS2* IR and *Th>NDUFB10* IR flies also exhibited prolonged morning sleep latency resembling that of mature control flies (**Figure 4J,K**), despite normal morning locomotor activity. Partial MCI inhibition, therefore, appears to preferentially disrupts characteristic features of juvenile sleep depth through increased wake-promoting drive.

### Depletion of reduced CoQ in DANs causes sleep loss

Our behavioral screen identified numerous hits in oxidative phosphorylation (OXPHOS) pathway components spanning MCI to MCV. Most OXPHOS perturbations, including manipulations directly targeting ATP synthase components, exhibited increased quiescence and locomotor suppression consistent with generalized energetic failure (**Figure 5A; Supplemental Figure 7A,B**). In contrast, only manipulations predicted to decrease the reduced coenzyme Q pool (CoQH_2_) caused robust sleep loss. In addition to the MCI perturbations described above, we identified *Th>sdhAF4* IR flies as short sleepers (**Figure 5A, B**). SDHAF4 is an assembly factor for Succinate Dehydrogenase/Mitochondrial Complex II(32), which oxidizes the TCA cycle intermediate succinate and transfers electrons to CoQ. Both RNAi lines comparably depleted *sdhAF4* mRNA (**Supplemental Figure 7C**), supporting the specificity of the phenotype. In juvenile flies, *sdhAF4* knockdown in DANs caused daytime sleep loss accompanied by elevated P(wake) (**Supplemental Figure 7D,E**), closely resembling the phenotype observed following partial MCI inhibition. Thus, perturbations predicted to deplete CoQH2 selectively (**Figure 5B)** promote wakefulness, whereas broader disruption of oxidative phosphorylation primarily suppresses locomotor activity.

**Fig 5:**
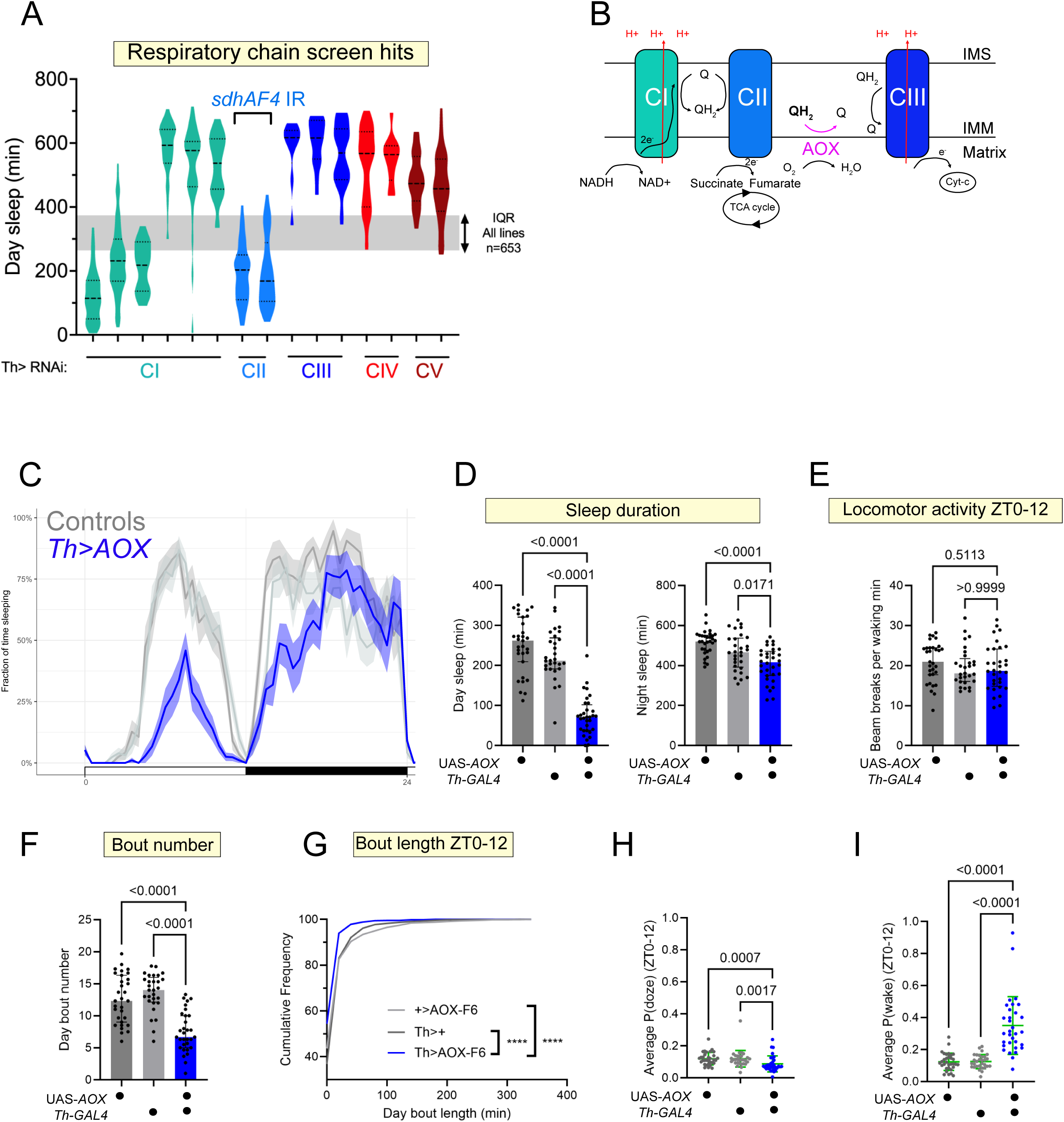
The CoQ redox state of dopaminergic neurons influences sleep. (A) Summary of daytime sleep phenotypes for all OXPHOS pathway screen hits. Grey shading reflects the interquartile range (IQR, the middle 50% of phenotypes) for all RNAi lines tested during the screen. While targeting any part of the respiratory chain can cause increased quiescence (due to reduced locomotor defects), only manipulations upstream of the CoQ pool (MCI and MCII) caused short sleep. (B) Schematic of respiratory complexes affecting the CoQ redox state. Reduced MCI and MCII activity both cause depletion of CoQH_2_. (C) Electron transfer at MCI can run in reverse, using electrons from CoQH_2_ to regenerate NADH at the expense of the PMV. The Alternative Oxidase (AOX) can directly deplete CoQH_2_ by transferring its reducing equivalents to oxygen. (D) Sleep per 30 min windows (average with 95% confidence interval shaded) of *Th>AOX* mature adult flies (blue) and genetic controls (grey). Depletion of CoQH2 was strongly wake-promoting, particularly during the ZT0-12. (E) Sleep duration during day (ZT0-12) and night (ZT12-24) for *Th>AOX* flies and genetic controls. Sleep loss was most pronounced during ZT0-12. (F) Waking locomotor activity during ZT0-12 for *Th>AOX* flies and genetic controls. Sleep loss was not associated with hyperactivity. (G) Number of sleep bouts for *Th>AOX* flies and genetic controls. AOX-overexpressing flies showed reduced sleep initiation. (H) Cumulative frequency plot of the durations of daytime sleep bouts for *Th>AOX* flies and genetic controls. AOX- overexpressing flies showed significantly reduced sleep bout duration. (**** p<0.0001. Kruskal-Wallis test with Dunn’s multiple testing comparisons) (I) P(doze) and (J) P(wake) averaged across ZT0-12 for *Th>AOX* flies and genetic controls. AOX overexpression affected both P(doze) and P(wake), but the effect on P(wake) was largest, indicating sleep loss in *Th>AOX* flies was driven largely by reduced sleep depth. P values computed using Kruskal-Wallis tests with Dunn’s multiple-testing comparisons.

To manipulate CoQ redox state independently of respiratory chain assembly, we expressed the Alternative Oxidase (AOX) from *Ciona intestinalis*, which oxidizes CoQH_2_ by transferring electrons directly to oxygen (**Figure 5B**)(33). Adult *Th>AOX* flies showed a pronounced daytime and early nighttime sleep loss driven by reductions in both sleep bout number and bout length (**Figure 5C, D, F, G**). Locomotor activity was unaffected (**Figure 5E**), indicating that sleep loss was not secondary to generalized hyperactivity.

Although both P(doze) and P(wake) were altered by AOX overexpression (**Figure 5H, I**), the most pronounced effect was an increase P(wake), indicative of reduced sleep depth and enhanced wake-promoting drive. Importantly, sleep loss was reproduced using two independent UAS-AOX insertions (**Supplemental Figure 7F-J**), ruling out insertion-site-specific effects. Together, these findings support a model in which maintenance of reduced CoQ levels within dopaminergic mitochondria restrains wake-promoting dopaminergic activity to promote sleep.

### Strong versus partial MCI LOF cause distinct metabolic phenotypes

The contrasting behavioral and cellular phenotypes associated with strong versus partial MCI LOF strongly suggested distinct metabolic phenotypes underpin them. To test this hypothesis, we expressed *NDUFB10* or *NDUFV1* interfering RNAs ubiquitously using the hormone-inducible *daughterless* (*da*) Geneswitch-GAL4 driver(34) and purified mitochondria to measure electron transport chain enzyme activity. In parallel, we also quantified the levels of NAD+, NADH, and the NAD/NADH ratio. Following eclosion, adult *da*GS-GAL4*>+, da*GS-GAL4>*NDUFB10* IR, and *da*GS- GAL4>*NDUFV10* IR flies were shifted onto 0.5 mM RU486-containing food for 5 days to induce RNAi expression. We observed a ∼56% reduction in MCI activity (**Figure 6A**) and no effect on MCII activity in *da*GS-GAL4>*NDUFV1* IR flies expressing the hypoactive RNAi line (**Figure 6B)**. By contrast, these measures of electron transfer activity were unaffected in *da*GS-GAL4>*NUDFB10* IR flies (Figure 6A,B). Thus, though both RNAi lines target essential MCI subunits (see Table 1 and Supplementary Figure 3), they have disparate effects on MCI enzyme function.

**Figure 6:**
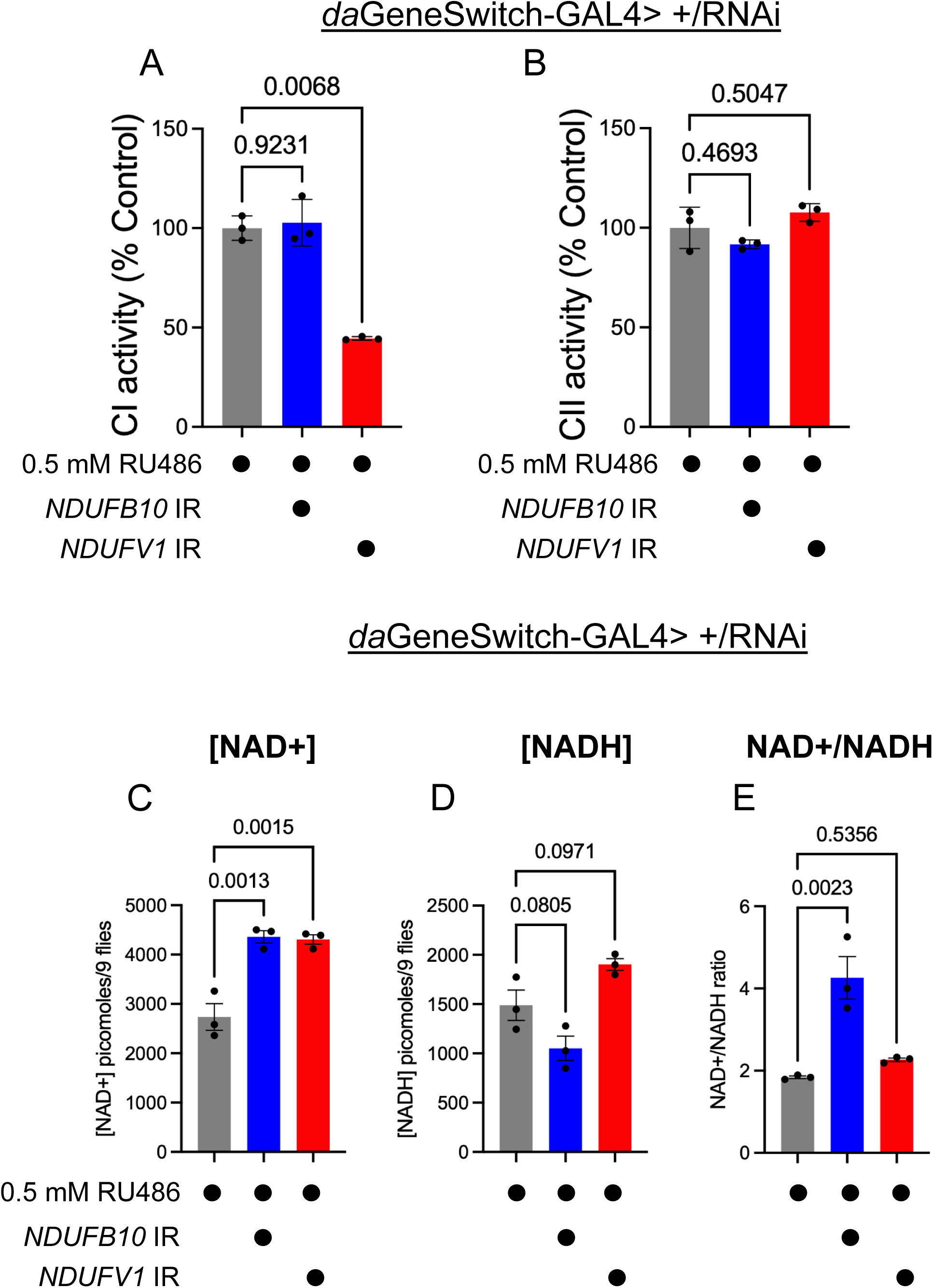
Strong versus partial MCI LOF cause distinct metabolic phenotypes. A) Measurements of MCI-dependent coenzyme Q reductase activity in crude mitochondrial fractions prepared from daGS-GAL4>+ (grey), daGS-GAL4>NDUFB10 IR (blue), and daGS-GAL4>NDUFV1 IR (red) flies shifted as adults onto standard fly food supplemented with 0.5 mM RU486 for 5 days. NDUFV1 knock-down reduced MCI activity to 56% of controls, while NDUFB10 knock-down had no effect on MCI activity. (B) Measurements of MCII-mediated coenzyme Q reductase activity in crude mitochondrial fractions of the same genotypes. Neither MCI perturbation affected mCII activity. P values computed using Browth-Forsythe and Welch ANOVA tests. (C-E) Measurements of NAD+ (C), NADH (D), and the NAD+/NADH ratio (E) in the same genotypes and same RU486 condition as in Panels A and B. NDUFB10 knock-down caused a pronounced increase in the NAD+/NADH ratio not observed with NDUFV1 knock-down. P values computed using ordinary one-way ANOVAs.

The enzyme activity assays described above measure electron transfer *in vitro* from exogenously supplied NADH to a synthetic and modified CoQ analog (CoQ1). Thus, we wondered whether *da*GS-GAL4>*NUDFB10* IR flies might have a subtler defect in MCI function that affects endogenous NAD redox balance, even if the MCI holoenzyme can convert synthetic substrates. Notably, d*a*GS-GAL4>*NDUFB10* IR flies exhibited a pronounced increase in their NAD+/NADH ratio (**Figure 6C-E**), indicating *NDUFB10* knock-down attenuates a process that normally exerts a strong downward effect on the NAD+/NADH ratio. By contrast, *da*GS-GAL4>*NDUFV1* IR flies exhibited elevations in both NAD+ and NADH levels, making their NAD+/NADH ratio unchanged relative to the control (**Figure 6C-E**). These experiments further indicate how strong versus partial MCI LOF cause metabolically distinct phenotypes. In addition to the forward electron transfer from NADH to CoQ (FET), electrons can also move in reverse through MCI, when the pool of reduced CoQ is abundant and the mitochondrial membrane potential strong. Reverse electron transfer (RET) regenerates NADH at the expense of NAD+, thereby driving down the NAD+/NADH ratio. Thus, inhibiting RET should increase the NAD+/NADH ratio, as shown previously for partial MCI inhibition (REF). Our data strongly suggest that the sleep loss phenotype is caused not by a defect in FET but by a more selective inhibition of RET.

### Coenzyme Q redox balance regulates juvenile DAN activity and influences later dopaminergic vulnerability

Because multiple independent genetic perturbations predicted to alter CoQ redox balance in DANs caused sleep loss, and because increased dopaminergic signaling promotes wakefulness(35), we hypothesized that RET modulates dopaminergic neuron activity. To test this idea, we measured cytosolic Ca2+ levels in juvenile DANs expressing UAS-*NDUFS2* IR. During the daytime period associated with elevated P(wake) (ZT4-8), *NDUFS2-*deficient DANs exhibited increased steady-state Ca2+ levels relative to controls (**Figure 7A,B**). In contrast, severe MCI disruption using *Th>NDUFV1* IR reduced in cytosolic Ca2+ levels (see **Figure 2I**), consistent with the profound locomotor hypoactivity observed under conditions of strong MCI impairment. Importantly, AOX expression in juvenile PPL1 DANs similarly increased in cytosolic Ca2+ levels (**Figure 7C, D**).

**Figure 7:**
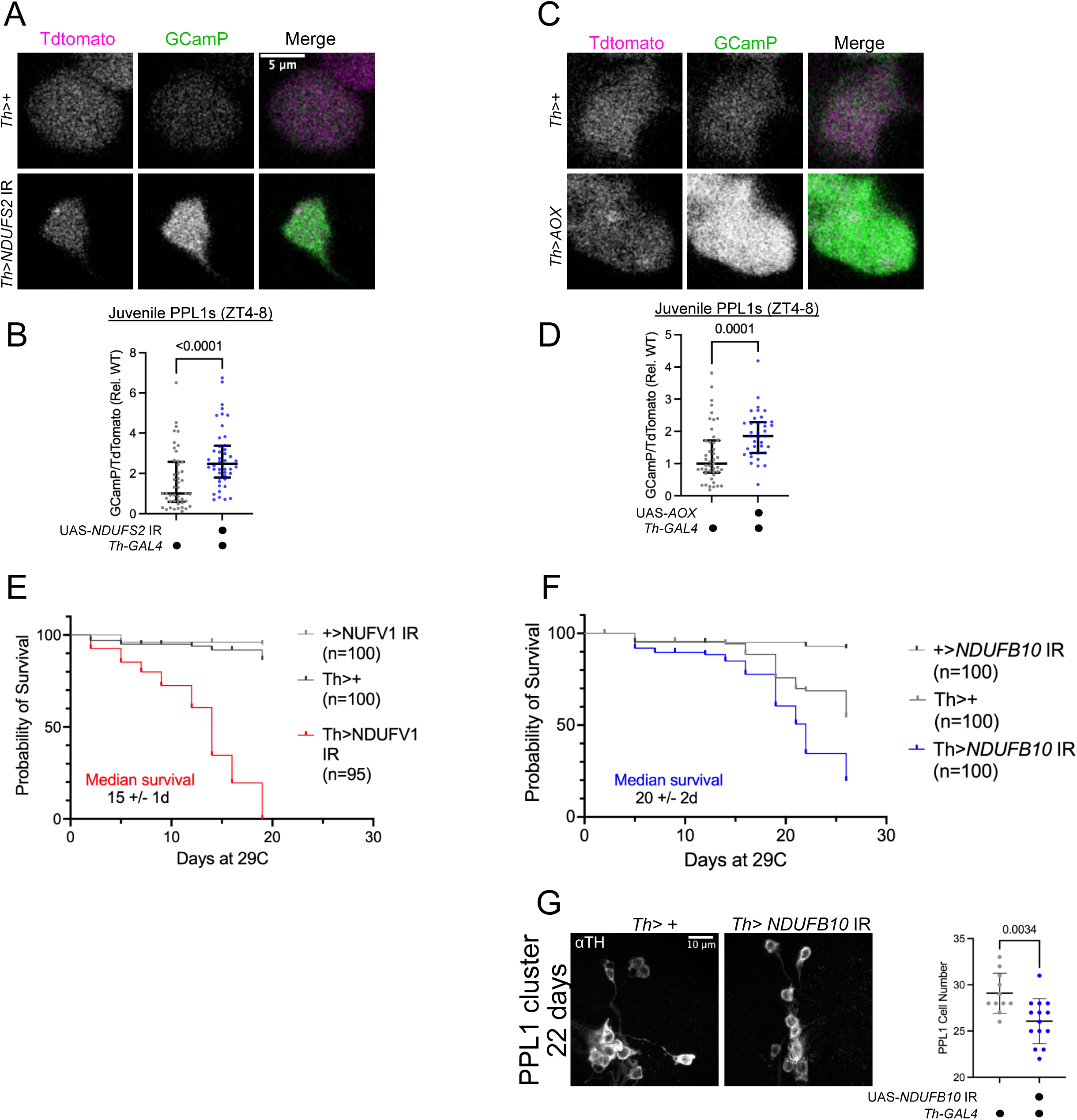
Depletion of UQH2 increases dopaminergic neuron activity. A) *Ex vivo* imaging of PPL1 DANs from *Th-LexA>TdTomato-2A-GCamP6s; Th-GAL4>+ attP2* (top) and *Th-LexA>TdTomato-2A-GCamP6s; Th-GAL4>NDUFS2* IR (bottom) juvenile brains. TdTomato is shown in magenta and GCamP in green. (B) Quantification of the GCamP/TdTomato ratios (median and interquartile range) in the juvenile PPL1 DAN cell bodies during ZT4-8 (n=6 female brains per condition from 2 independent experiments). Ratios are normalized to the median of the control brains imaged in parallel. Partial MCI inhibition caused an increase in cytosolic Ca^++^ levels, in contrast to the reduction seen with severe MCI RNAi. P value computed by a Mann-Whitney t-test. (C) *Ex vivo* imaging of PPL1 DANs from *Th-LexA>TdTomato-2A-GCamP6s; Th-GAL4>+* (top) and *Th-LexA>TdTomato-2A-GCamP6s; Th-GAL4>AOX-F6* (bottom) juvenile brains. (D) Quantification of GCamP/TdTomato ratios (median and interquartile range) in juvenile PPL1 DAN cell bodies during ZT4-7 (n=4 female brains per condition from 2 independent experiments). Ratios are normalized to the median of the control brains imaged in parallel. AOX overexpression elevated cytosolic Ca^++^ levels in DANs. P value computed by a Mann-Whitney t-test.

Severe MCI disruption caused early dopaminergic dysfunction and locomotor impairment (see **Figure 2**). Consistent with the well-established link between MCI dysfunction and dopaminergic neurodegeneration(36), *Th>NDUFV1* IR flies also exhibited markedly shortened lifespan (**Figure 7E**). These findings raise the possibility that prolonged partial MCI perturbation might similarly compromise neuronal integrity over time. Indeed, flies with partial MCI inhibition in DANs (*Th>NDUFB10* IR) also died prematurely, but with delayed onset relative to severe MCI disruption (**Figure 7F**).

Moreover, aged *Th>NDUFB10* IR flies exhibited a loss of PPL1 dopaminergic neurons compared to genetic controls (**Figure 7G**). Thus, partial MCI inhibition is sufficient to drive age-dependent dopaminergic vulnerability despite initially producing a wake-promoting rather than hypofunctional phenotype.

## Discussion

Elevated sleep during early life supports critical processes of brain development, yet the cellular mechanisms that maintain the juvenile sleep state remain poorly understood. Here, we identify mitochondrial Complex I and coenzyme Q redox state in dopaminergic neurons as regulators of sleep drive. Transcriptomic profiling revealed enrichment of oxidative phosphorylation and redox-related genes in juvenile dopaminergic neurons, suggesting that early life is associated with a distinct mitochondrial state. While severe disruption of MCI caused dopaminergic hypofunction and locomotor impairment, partial perturbations predicted to deplete the reduced coenzyme Q pool increased dopaminergic activity and decreased sleep. Because reduced CoQ provides the electron donor for reverse electron transfer (RET) at MCI, our results support a model in which RET-related mitochondrial signaling physiologically restrains dopaminergic activity. Our biochemical analyses of MCI enzyme activity and NAD redox balance under the different RNAi conditions described above are consistent with this interpretation. Although this mechanism is not exclusive to juvenile flies, its effects are especially pronounced during early life, when elevated sleep drive depends on suppression of dopaminergic activity.

Disturbed mitochondrial energetics is a primary cause of dopaminergic degeneration leading to Parkinsonism. In particular, both environmental and genetic drivers of the motor symptoms of Parkinsonism converge on reduced MCI activity(36). There is mounting evidence that sleep disturbances may function as early harbingers, or even drivers, of later neurodegeneration(37). We identified a surprising consequence of partial MCI disruption in DANs. Rather than producing an intermediate locomotor phenotype between wild-type and severe MCI LOF, partial MCI inhibition caused sleep loss accompanied by increased sleep-to-wake transitions and elevated dopaminergic activity. These findings suggest that mitochondrial dysfunction may not progress monotonically from normal neuronal function to hypoactivity, but instead pass through an early hyperexcitable state characterized by augmented DA arousal signaling.

Interestingly, dopaminergic-specific ablation of MCI activity in mice similarly causes sleep fragmentation before the onset of motor symptoms and neuron loss(38). In our experiments, partial MCI inhibition ultimately resulted in shortened lifespan and age-dependent dopaminergic neuron loss despite the absence of early signs of dopaminergic hypofunction. Together, these observations raise the possibility that early sleep disturbances driven by altered mitochondrial signaling may represent sensitive indicators of emerging dopaminergic vulnerability.Importantly, augmented DAN activity cannot be explained merely by a collapse of mitochondrial function or ATP depletion.

While severe MCI disruption caused mitochondrial swelling, locomotor hypoactivity, and reduced calcium activity in DANs, manipulations predicted to deplete the reduced coenzyme Q pool (CoQH2) selectively promoted wakefulness and increased dopaminergic activity. This effect was reproduced using multiple independent manipulations, including partial inhibition of MCI, disruption of MCII assembly, and expression of Alternative Oxidase (AOX), indicating that altered CoQ redox balance rather than generalized respiratory failure underlies the wake-promoting phenotype. Because CoQH2 is the source of electrons for reverse electron transfer (RET) at MCI, our results suggest that dopaminergic MCI maintains a basal level of RET to *limit* neuronal activity. Interestingly, mitochondria isolated from juvenile flies exhibited greater electron transfer capacity to coenzyme Q than mitochondria from mature adults. While previous work has shown that the pathological effects of RET at MCI (e.g. oxidative damage) increase with advanced age(18), we speculate that the more efficient electron transfer to the juvenile CoQ pool allow RET to exert strong physiological effects in juvenile periods. In this way, subtle defects in mitochondrial redox signaling could disrupt sleep well before the onset of overt neurodegeneration.

Prodromal changes to mitochondrial function in Parkinson’s disease may first manifest as changes to RET as even small perturbations can influence this route of electron transport. While forward electron transfer is thermodynamically favorable under broad physiological conditions, RET is tightly coupled to a specific set of metabolic parameters (high membrane potential, abundant ubiquinol, and high NAD+/NADH ratio) and likely operates within a narrow threshold of feasibility. Partial MCI inhibition may cause quantitative reductions in these parameters that shift mitochondrial conditions outside of this threshold. In this way, neuronal processes downstream of RET-dependent redox signaling may be the first to fail before overt neurodegeneration caused by mitochondrial dysfunction.

How might alterations in CoQ/CoQH₂ redox state increase dopaminergic neuron activity? One possibility is that partial inhibition of MCI and RET produces modest changes in redox signaling that enhance neuronal excitability. In neurons, low-level increases in reactive oxygen species (ROS) can potentiate activity by modulating voltage-gated ion channels, intracellular Ca²⁺ handling, or neurotransmitter release(39–41), whereas more severe or sustained redox stress leads to mitochondrial damage and neuronal hypoactivity(41). Shifts in mitochondrial membrane potential, NAD⁺/NADH ratios, or ATP availability at presynaptic sites could increase release probability or excitability(42, 43). Such mechanisms may be particularly relevant in juvenile dopaminergic neurons, which exhibit elevated metabolic gene expression and may rely on tightly regulated redox states to restrain wake-promoting output.

These findings add to a growing body of work implicating mitochondria as key regulators of sleep need. Previous studies have linked changes in homeostatic sleep pressure to changes in neuronal mitochondrial redox signaling, including within defined sleep-promoting neuronal populations(44–46). Our work identifies a mechanism in which subtle perturbations of MCI and CoQ redox state selectively modulate dopaminergic output to regulate sleep drive and arousal state. By dissociating mitochondrial redox signaling from ATP depletion or respiratory collapse, our results support a model in which coenzyme Q redox balance functions as a signaling node within wake-promoting circuits. This regulation is most apparent during juvenile periods, when elevated sleep drive depends on suppression of dopaminergic arousal. More broadly, these findings suggest that mitochondrial mechanisms contributing to long-term dopaminergic resilience may also help maintain early-life sleep states, linking sleep early in life to later neuronal vulnerability.

## Materials and Methods

### Cell sorting and RNAseq

Juvenile (0-1 day old) and mature (6-9 day old) *Th*>TdTomato brains were dissected in ice-cold Schneider’s medium. Brains were first enzymatically dissociated in Papain (100U/ml) suspended in 1X PBS and supplemented with 0.18U/ml Liberase^TM^ cocktail (Roche). Dissociation was carried out for 50 min at room temperature with nutating.

Dissociated brains were rinsed three times in cold Schneider’s to stop dissociation. Schneider’s medium was replaced by 2% Bovine Serum Albumin in 1XPBS. Mechanical trituration was performed by first pipetting the brains up and down 100% in P200 pipette tips pre-coated with fetal bovine serum; then, dissociated tissue was passed 20 times through a 3cc gauge needle pre-coated with fetal bovine serum. The suspension was then transferred to FACS tubes kept on ice. DAPI was added to label dead cells. In parallel, +>TdTomato brains were treated in parallel to gate TdTomato signal during FACS sorting. Sorting was performed at the Penn Cytomics core using an Aria cytometer equipped with a 450 nm UV and 582 nm lasers and a 100 um nozzle.

FACSDiva (v8.0.1) was used to analyze data. Dopaminergic cells were gated by low DAPI and high TdTomato signal. Cells were sorted directly into a lysis buffer for the SMART-seq HT kit (Takara Biosciences) and flash frozen on dry ice. Library preparation and RNA sequencing was performed using the Illumina NGS platform by Admera Health (South Plainfield NJ, USA) to obtain paired-end 150 bp reads. Trim Galore (v0.4.4_dev) and Cutadapt (v2.4) were used to remove Adapter Sequences from the reads. Salmon (v1.10.1) was used to quantify transcript read counts against transcripts from v110 of the Ensembl Drosophila Melanogaster annotation. Using R (v4.3.1), the Salmon output was summarized to gene-level quantifications using the tximeta package (v1.18.3). DESeq2 (v1.40.2) was used to perform differential expression analysis between the juvenile and mature samples on the unnormalized read counts.

### RNAi screen

Virgin *Th-GAL4* females were mated to males from the Transgenic RNAi Project lines carrying UAS-dsRNAs or UAS-shRNAs carried in VALIUM10, VALIUM20, or VALIUM22 vectors(47, 48) . All RNAi lines were obtained from the Bloomington Drosophila Stock Center (Bloomington, IN, USA) 16 mated UAS-dsRNA/+; *Th-GAL4/+* or Th-GAL4/UAS- dsRNA females were transferred into glass tubes containing 5% sucrose in 2% agar as a food source and loaded into DAM2 activity monitors (Trikinetics, Princeton, MA, USA). Sleep was assayed under 12h:12h light:dark cycles in 6-9 day old flies. Each RNAi line was tested in 2 independent biological experiments run at separate times in the same incubator. Average sleep duration for ZT0-12 was used to identify short- and long- sleepers.

### Sleep assays

Flies were maintained in a 12h:12h light:dark cycle throughout their lifecycle. Virgin females were used for juvenile (1 day post eclosion) and mature adult (5-7 days post eclosion) sleep assays. Flies were anesthetized using CO2 and transferred into glass tubes containing 5% sucrose in 2% agar as a food source. DAM5H activity monitors (Trikinetics, Princeton, MA, USA) were used to track locomotor activity in 12h:12h light:dark cycles at 25C and ∼60% relative humidity using Percival incubators. All sleep assays were performed in the same incubator throughout. Activity count data were analyzed using the Rhethomics pipeline(49) in R for most sleep metrics (duration, bout number, bout length, activity index). Pwake and Pdoze were calculated using the Sleep and Circadian Matlab Analysis Program (SCAMP)(50). In all analyses, bouts of inactivity >/= 5 min were used as the standard definition of sleep. Sleep latency times were calculated by manually scoring the time to the first sleep bout (inactivity for >/= 5 min) directly from raw activity counts.

### Survival Analysis

100 Mated females of each genotype were housed in groups of 20 at 29C under a 12h:12h light:dark cycle. Every 2-3 days, the number of dead flies was counted and the living flies were flipped onto fresh food. Kaplan-Meier survival analyses were performed using Prism software. Each survival experiment was repeated at least twice to measure an average median survival time +/- standard deviation.

### Fly husbandry

Fly crosses were reared on standard laboratory media (LabExpress) at 25C on a 12h:12h light:dark cycle under ∼60% relative humidity. Below are listed fly stocks used in this paper.

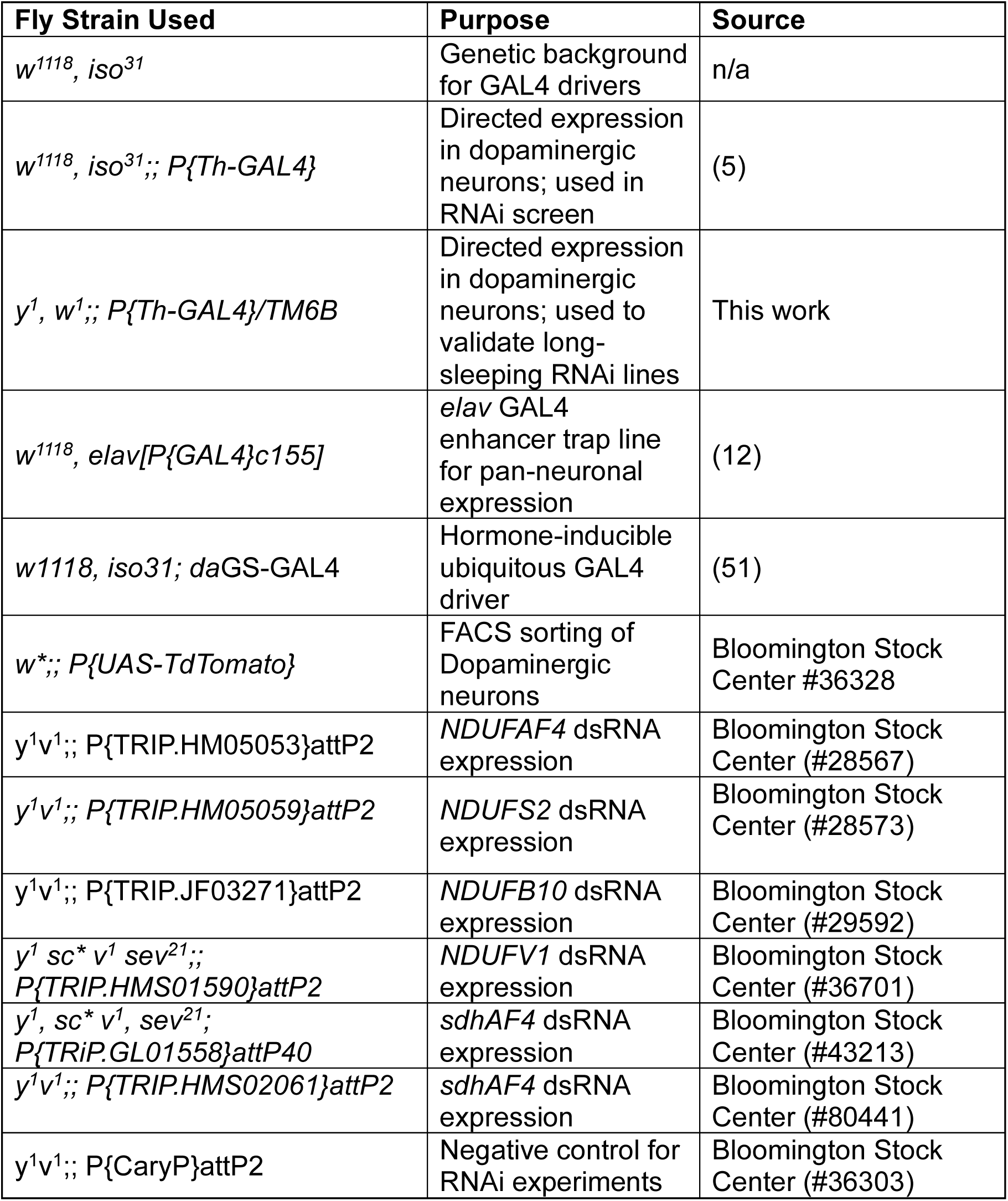
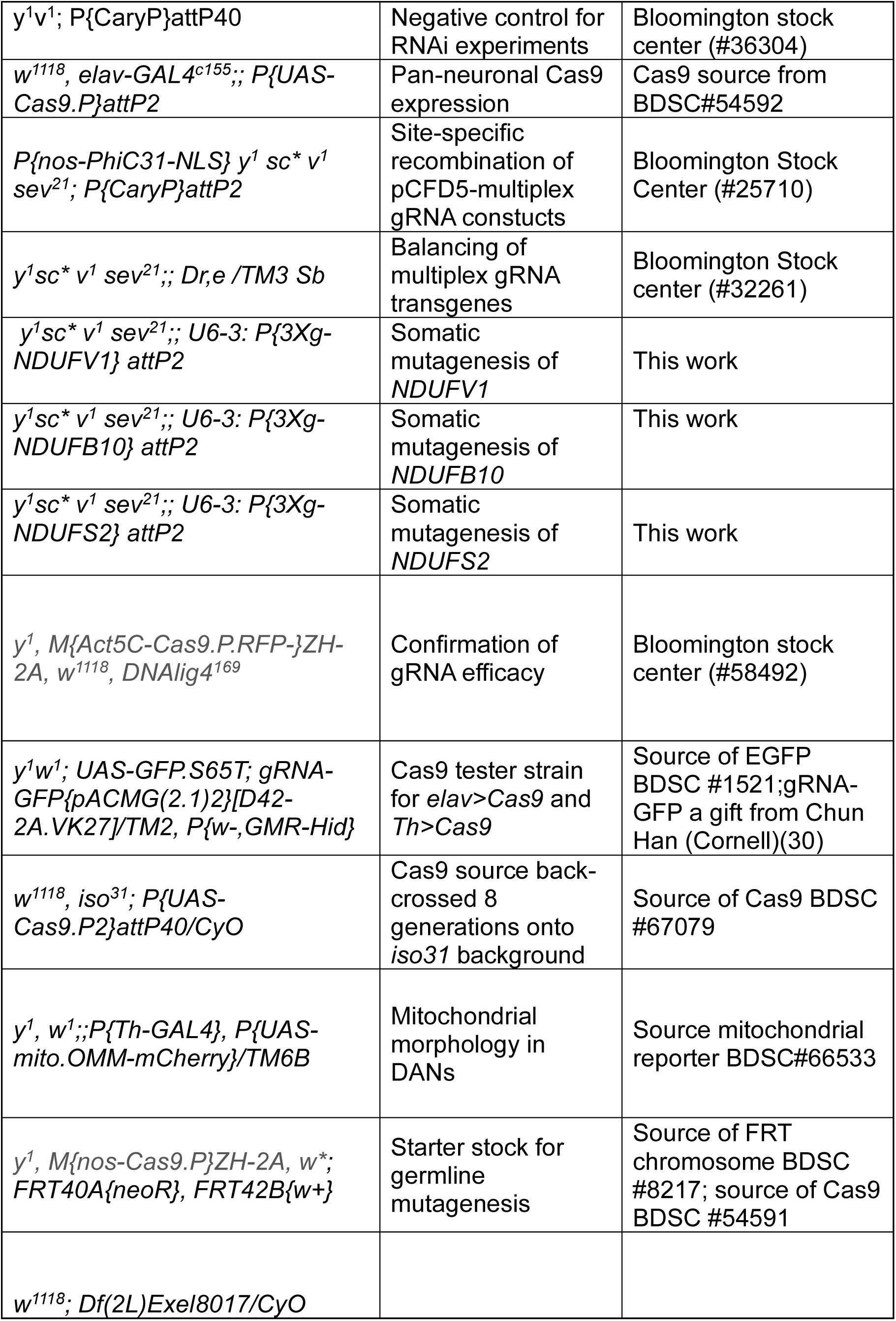
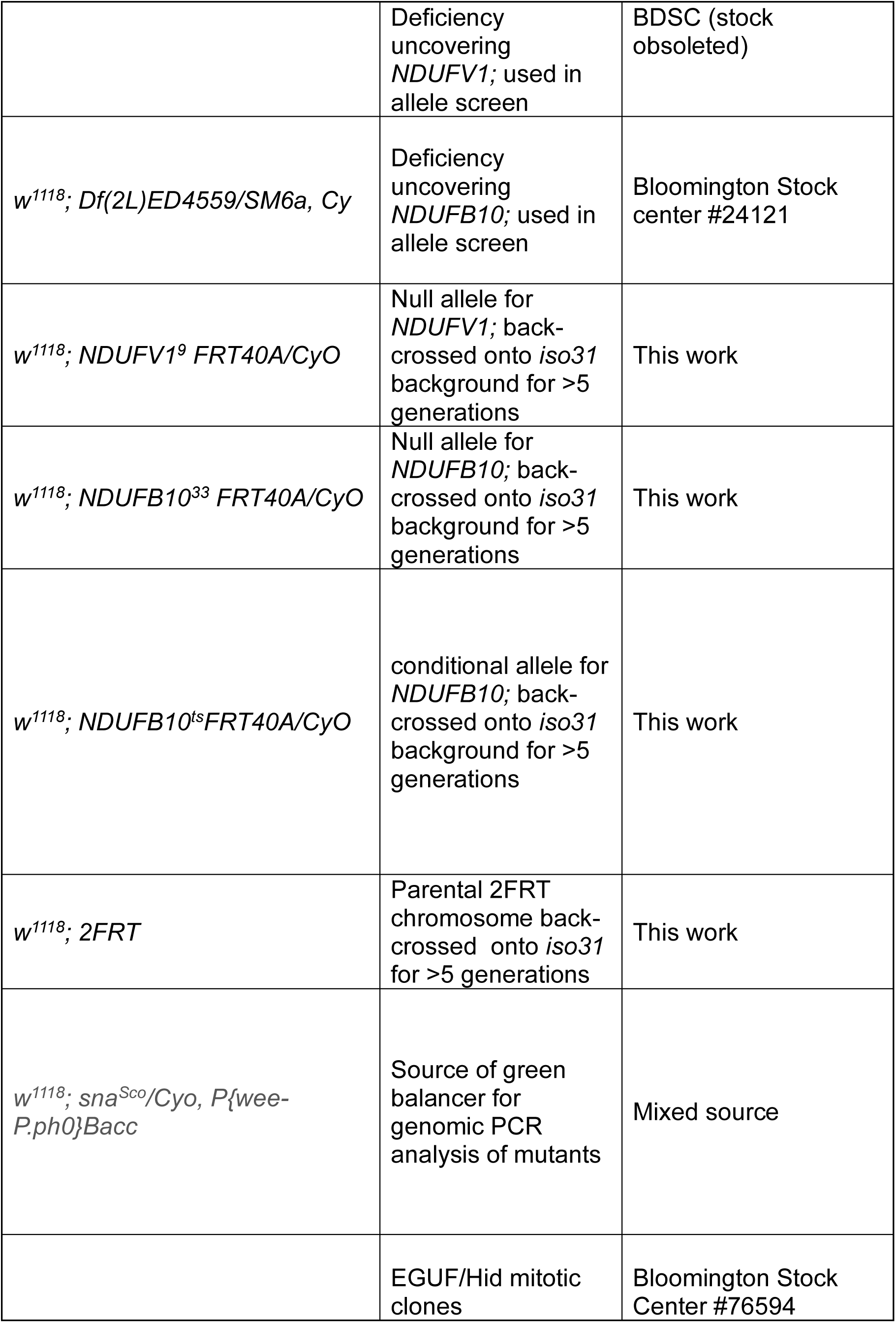
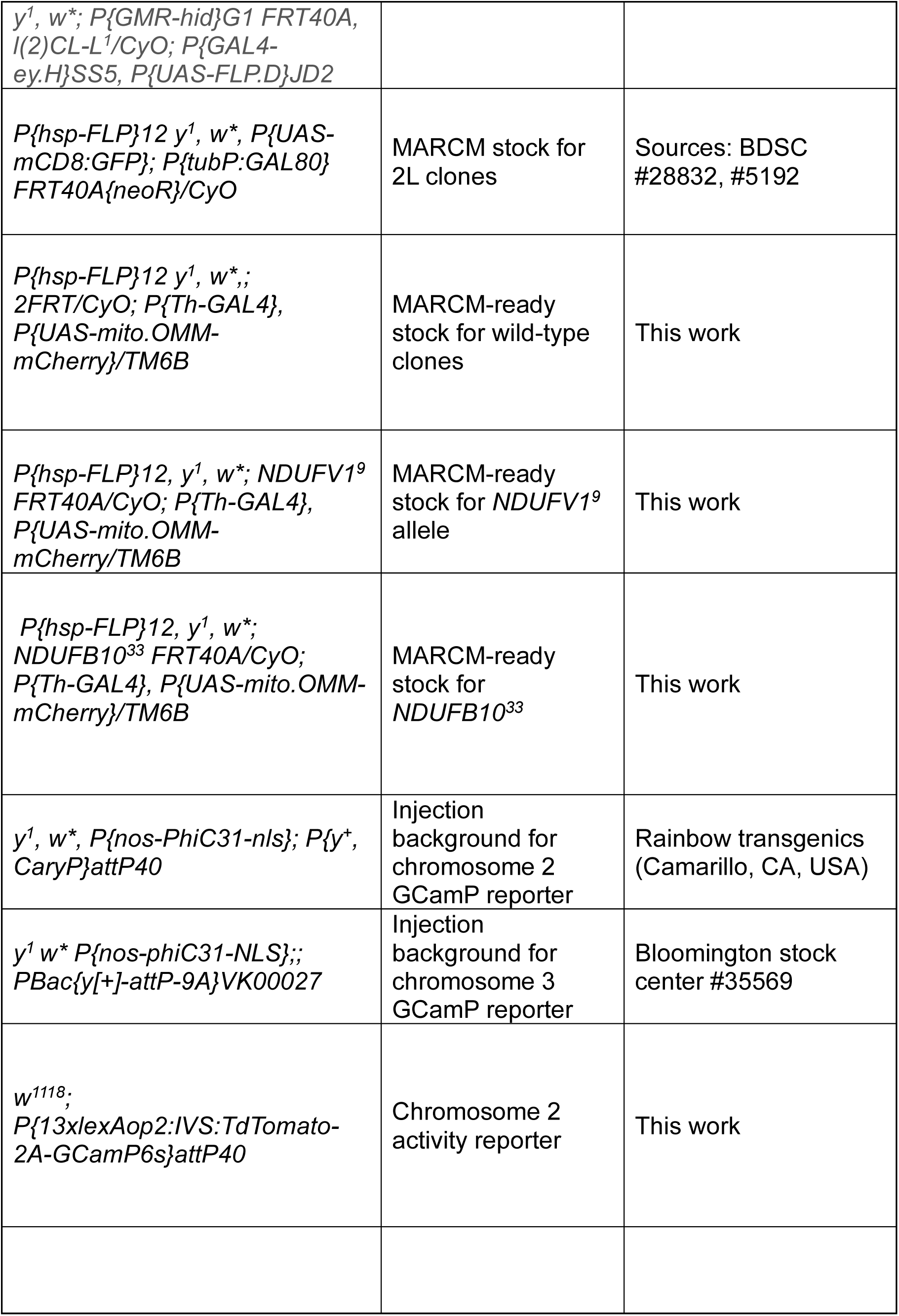
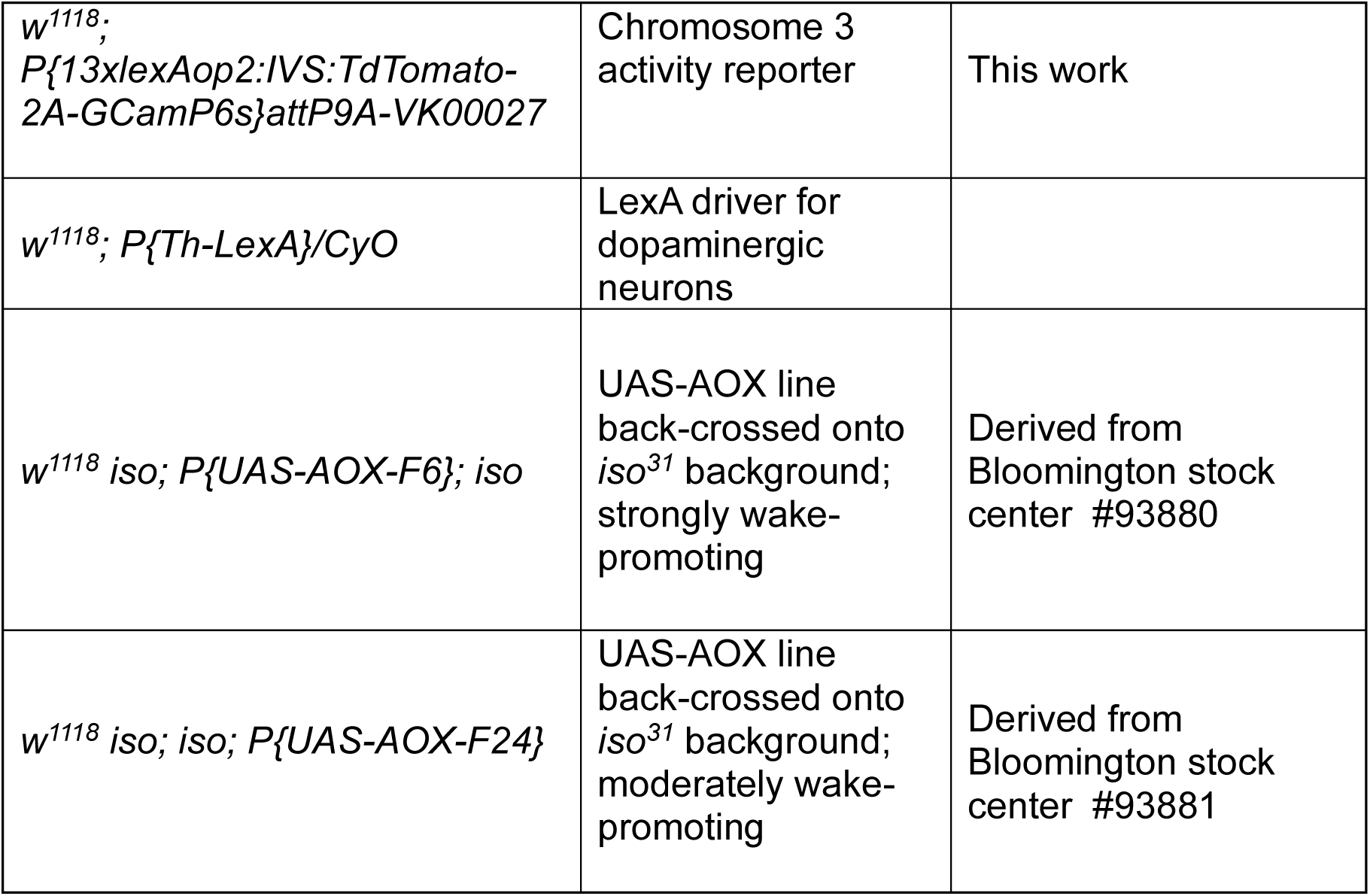

### Transgenic guide RNA construction

The protocol of Port and colleagues was used to generate multiplex gRNA vectors for transgenesis(17). Using primers described in Supplemental Table 2, 3 PCR fragments (for each target gene) encoding tRNA-flanked exonic sgRNAs were amplified off a *pCFD5* plasmid template (Addgene #73914). Oligos were synthesized by Integrated DNA Technologies. The two largest oligos containing homology arms for Gibson assembly were PAGE-purified after synthesis to ensure correct sequence. sgRNA templates were generated using Q5 DNA Polymerase (New England Biolabs) using the following thermal cycling conditions: 98C for 30S-->32 cycles of 98C for 10, 61C (+0.5C/cycle) for 30s, 72C for 10s-->72C 2min. Gel-purified (ZymoClean Gel DNA Recovery Kit, Zymo Research) PCR amplicons were combined in a 5-fold molar excess with BbsI-linearized *pCFD5* in a Gibson assembly reaction (New England Biolabs). The *pCFD5* plasmid allows for the expression of multiple sgRNAs under the control of a single ubiquitous RNA Polymerase III promoter. Colony PCR using a GOTaq master mix (Sigma) was used to identify correctly assembled clones using the following thermal cycling parameters: 95C for 2 min--> 25 cycles of 95C for 30s, 54C (+0.5C/cycle) for 30s, 72C for 1 min-->72C for 5 min). Candidate clones were expanded in liquid culture supplemented with ampicillin (10 ug/ml) and extracted using QIAprep mini plasmid kits (Qiagen) for Sanger sequencing. See Supplemental Table 2 for primer information.

Sanger sequence-validated plasmid DNA was prepared using a Midi plasmid DNA kit (Qiagen) and injected into *y M{nos-Phi*C31*} v;; P{CaryP}attP2* by Rainbow Transgenics (Camarillo, CA, USA). Individual G0 flies were crossed to *y v;;Dr/TM3,Sb* (Bloomington stock #3261) to recover germline integrants. *vermillion+* G1s were back-crossed to recover a balanced line without the *M{nos-Phi*C31*}* transgene. To test cutting efficacy of the gRNAs, transgenic gRNA lines were crossed to an *Act5C-Cas9.P, DNAlig^4^* stock to induce somatic mutagenesis ubiquitously and in the presence or absence of DNA repair: neither *Act5C-Cas9, DNAlig^4^;;* U6-3:gRNA nor *Act5C-Cas9 DNAlig^+^;;* U6- 3:gRNA adults were recovered, reflecting the essential role for MCI function.

### Germline mutagenesis

An *FRT40A P{neoR}, FRT2A P{mini-w+}* (hereafter *2FRT*) chromosome was isogenized by standard crossing methods. Females from a M{*nos-*Cas9}; *2FRT* isogenic stock were mated with males carrying multiplex gRNA transgenes on chromosome 3.

The resultant *nos-Cas9/Y; 2FRT/+; gRNA/+* males were then mated to *w^1118^; Sco/CyO* females to recover *\* FRT40A* /Sco males carrying unique mutagenized chromosomes. Single males were mated to females carrying hemizygous chromosomesal deficiencies uncovering *NDUFB10* or *NDUFV1.* New recessive lethal alleles were identified by allelic non-complementation, and * FRT40A/CyO sibling flies were collected for a stock. Non-complementing lines were crossed to females from a yw; P{GMR-Hid} FRT40A l(2)CL-1; ey>FLP stock to induce mitotic recombination in the eye. Mutagenized chromosomes causing the most severe eye malformations were chosen for genomic sequencing.

Eggs from *\* FRT40A/CyO, weeP* in-crosses were collected on standard apple juice/agar plates for ∼16-18h at 25°C and aged for 24h at 25°C. GFP- larvae were picked under a fluorescence stereomicroscope and homogenized in 50 ul of squish buffer (10mM Tris, 25 mM NaCl, 1 mM EDTA supplemented with 200 ug/ml Proteinase K), followed by activation of the Proteinase K under these conditions: 30 min at 37C, 2 min at 95C. See Supplemental Table 2 for primers used for PCR amplification and Sanger sequencing of *NDUFB10* and *NDUFV1* genomic regions. Taq DNA Polymerase (New England Biolabs) was used in the following thermal cycling conditions: 95C for 3 min-->35 cycles of 95C for 30s, 51C for 30s, 72C for 1:15s-->72C for 5 min. Sequences from mutants were compared to those derived from the *w^1118^; 2FRT* stock . Alignments to the reference genome (r6.65, Flybase.org) were performed using SnapGene. Mutants of interest and the parental chromosome were back-crossed for >5 generations onto an isogenic *w^1118^* genetic background.

### Mitotic clone generation

Eggs of the genotype *UAS-mCD8:GFP, hsp70-FLP12; tubulinP-GAL80 FRT40A/ * FRT40A; Th-GAL4>mito.OMM:mCherry/+* were collected on standard fly food at 25°C for ∼18h and aged to 48-66h after egg-laying. FLP expression was induced using a 38°C heat shock for 30 min in a water bath. Brains from females 2-4 days old adults were dissected and fixed for immunostaining.

### Generation of lexAop:TdTomato-2A-GCamP6s reporter

A PspXI-TdTomato-2A-GCamP6s-XbaI fragment was PCR amplified from Addgene plasmid #130666 and subcloned into the PspXI and XbaI sites of the linearized backbone of Addgene plasmid #111548 using standard subcloning methods. This placed the TdTomato-2A-GCamP6s coding sequencing under the control of a 13xLexAop promoter. See Supplemental Table 2 for PCR and sequencing primers used. PhiC31 integrase was used to recombine the attB site on the plasmid backbone at chromosomal attP dock sites on chromosome 2 (attP40) and chromosome 3 (attP9A).

Germline transformants were selected by mini-w+ expression off the integrated transgene and stable lines were back-crossed onto a *w^1118^ iso^31^* background. Injections were performed by Rainbow Transgenics (Camarillo, CA, USA).

### Quantitative PCR

UAS-dsRNAs were expressed using the pan-neuronal *elav-*GAL4 driver and compared to a GAL4+/dsRNA- control of the same genetic background (e.g. *elav>+ attP2)*. ∼15 heads were used per genotype per biological triplicate. Total RNA was isolated from whole heads flash-frozen on dry ice and homogenized in Trizol (Invitrogen). For *NDUFV1* knock-down quantification, dissected brains from wandering third instar larvae (∼25 per replicate) were used. Briefly, nucleic acids were extracted in a 5:1 mixture of Trizol: chloroform. Following centrifugation (12,000*xg* at 4C, 15 min), the upper polar phase was subjected to isopropyl alcohol precipitation (1:1 aqueous phase: isopropyl alcohol, 20 ug glycogen). Nucleic acid pellets were washed in 75% ethanol, dried, and then resuspended in 30 ul DEPC-treated ddH20. DNAse digestion was performed using Turbo® DNase (ThermoFisher) using manufacturer’s protocol. Total RNA concentration was measured using a NanoDrop spectrophotometer. 1 ug of total RNA was used for first-strand cDNA synthesis (SuperScript RT II, ThermoFisher) primed with random hexamers (ThermoFisher). qPCR primers for each target cDNA were designed using NCBI Primer Blast(52). Oligos were synthesized by Integrated DNA Technologies and used at a concentration of 0.2uM. SYBR® green PCR master mix (Applied Biosystems) was used. PCR reactions were run using the BioRad CFX Opus 96 thermal cycler. The amplification efficiency of each primer pair and control primers (see Supplementary Table 1) was calculated from the slope of the linear regression of Ct values obtained from amplification off a 5-fold dilution series of wild-type whole-head cDNA, according to the formula *efficiency= 10^-(1/slope)^ -1*. qPCR primers amplifying single amplicons with >95% efficiency were used in calculating relative mRNA abundance. The ΔΔCt method was used to calculate relative mRNA abundance. Data were analyzed using Excel and statistics performed using GraphPad Prism software.

### Mitochondrial Complex I assays

Mitochondrial fractions were prepared from gently homogenized flies following the protocol of Groen and Windebank(53). ∼50 flies of mixed sex were collected per replicate per genotype. Mitochondria were isolated and resuspended on ice in 70 mM sucrose, 210 mM mannitol, 5 mM HEPES, 1 mM EGTA, and 0.5% bovine serum albumin, pH 7.5. Protein concentration was measred using a Pierce™ BCA assay kit (Thermo Scientific). Complex I and complex II enzyme activities were determined by the reduction of 2,6-DCPIP at 600 nm (ε600 = 21 mM−1 cm−1). The assay buffer for both complex I and II enzyme assays contained 25 mM KH2PO4, 5 mM MgCl2, 3 mg/mL bovine serum albumin (BSA), 25 µM ubiquinone Q1, 5 μM antimycin A at pH 7.4. The reactions were initiated by the addition of 150 µM NADH in the presence and absence of 5 μM rotenone, and the complex I rates were calculated after subtraction of the ROT- insensitive activity. The enzyme activity assay solution for complex II always contained 5 μM ROT to inhibit complex I, along with fly crude mitochondrial fractions. The reactions were started by the addition of 20 mM succinate. For RNAi induction of MCI RNAi lines, a 0.5 uM RU486 solution (in 200-proof ethanol) was diluted 100-fold in molten standard fly media. 1-3 day old adults were transferred onto RU486-containing food for 6d and flipped onto fresh food every 2-3 days uner 12h:12h light:dark conditions at 25C. Mitochondria were harvested from whole flies (∼50-60 females per replicate per genotype).

### NAD/NADH measurements

UAS-RNAi lines were induced by feeding adult flies daGS-GAL4>RNAi flies 0.5 mM RU486 added to standard fly food for 5d. Mated female flies were transferred onto fresh RU486 food every 2 days and kept at 25C under 12h:12h light:dark conditions. 9 carcasses were used for each nucleotide extraction for each biological replicate and each genotype. Flash-frozen headless flies were homogenized in either 0.6N perchloric acid or 0.375N potassium hydroxide/50% ethanol for NAD+ and NADH, respectively, using a handheld homogenizer. Homogenized extracts were centrifuged at 10,000 x g for 10 minutes at 4 °C. The resulting supernatants were diluted 1:100 or 1:50 for NAD+ and NADH, respectively, in ice-cold 100 mM sodium phosphate buffer (pH 8.0), and NAD+ was measured by an enzymatic cycling assay modified from the protocol of Graeff and Lee (54) 5 uL of extracted sample or standard was mixed with 95 uL of cycling mix and NAD+ concentration was calculated depending on the rate of resorufin accumulation, which was measured as fluorescence at excitation at 544 nm and emission at 590 nm as described in(55)

### Immunostaining

Adult brains were dissected in cold 1X PBS and fixed in 4% paraformaldehyde (Electron Microcsopy Sciences) for 20 min. Tissue was permeabilized in 0.3% Triton X-100 in 1X PBS (PBST) for 20 min. All steps performed at room temperature with nutating. Blocking was performed in 5% goat serum in PBST for >1h at room temperature or overnight at 4C. Primary antibodies were diluted in PBST with 5% goat serum and incubated with nutating at 4C overnight. Secondary antibodies were diluted 1000-fold in PBST and incubated with brains for 2h at room temperature or overnight at 4C. Between antibody incubations, brains were washed in PBST at room temperature with nutating for 20 min. Before mounting, PBST was replacted with 1X PBS and the brains were mounted on slides with a 2:1 mixture of SloFade mounting media: 1X PBS. Imaging spacers were placed between the slide and coverslip. Primary antibodies used were: Chicken anti-GFP (Invitrogen) at 1:1000, Rabbit anti-mCherry at 1:100 (GeneTex), rabbit anti-Tyrosine Hydroxylase at 1:100 (BD Biosciences), and nc82 at 1:100 (Developmental Studies Hybridoma Bank). Secondary antibodies (Invitrogen) used were: Goat anti-Chicken Alexa 488, Donkey anti-Rabbit Alexa 555, and Donkey anti-mouse Alexa 647.

### Microscopy

Imaging was performed on a Leica SP6 laser scanning confocal microscope (Leica Microsystems) equipped with 488 nm, 514 nm, and 613 nm lasers. 512 x 512 images z-stacks (1.5-2um step sizes) were collected by bidirectional linear scanning with 2x line averaging. For fixed and stained samples, scanning was performed at 400Hz. Where quantitation of intensity was performed across conditions, laser power and gain were kept constant across conditions. For qualitative imaging, power and gain were adjusted to bring pixel intensity just below saturation. For *ex vivo* GCamP imaging, brains were rapidly dissected during ZT1-3 and mounted in a drop of cold Artificial Hemolymph (108 mM NaCl, 5 mM KCl, 2 mM CaCl2, 8.2 mM MgCl2, 4 mM NAHCO3, 1 mM NaH2PO4, 5 mM Trehalose, 10 mM Sucrose, 5 mM HEPES, adjusted pH 7.5) caudal side up on poly-l-lysine slides with imaging spacers (Grace Bio-Labs, ref#654008) and a coverslip. Laser scanning was performed at 8000Hz using resonant scanning using 488 nm and 514 nm lasers. Only the PPL1 DAN cell bodies closest to the objective were imaged to minimize light scattering. Linear bidirectional scanning with 2x averaging and a 72 ns pixel dwell time was used throughout all experiments. Laser power and gain were kept constant during imaging sessions. Single optical slices containing the majority of cell body signal were used for acquiring images every 30s for ∼2.5-5 min to assess the stability of GCamP fluorescence. GCamP6S fluorescence never varied across time- lapse imaging, but the same frame (t=3) was used for all image quantitation. A 3x3 smoothing filter in ImageJ was applied to the EGFP and TdTomato channels before calculating the background-subtracted mean intensity of each cell body. The ratio of mean GCamP/TdTomato intensities was calculated for each cell body. All mean intensity measurements were made using ImageJ.

## Acknowledgments

We thank members of the Kayser Lab and of the Penn Chronobiology and Sleep Institute for helpful discussion and input.

## Funding

Simons Foundation Autism Research Initiative, Life Science Research Foundation (JBR)

NIH R01NS120979 (MSK)

NIH R35NS137329 (MSK)

Burroughs Wellcome Career Award for Medical Scientists (MSK)

NIH RO1DK098656 (JAB)

## Author contributions

Conceptualization: JBR, MSK

Investigation: All authors

Writing – Original Draft: JBR, MSK

Writing – Review and Editing: All authors

Project Supervision and Funding: MSK

## Data and materials availability

All data needed to evaluate the conclusions in the paper are present in the paper and/or the Supplementary Materials. Raw RNA sequencing data can be accessed at https://www.ncbi.nlm.nih.gov/geo/query/acc.cgi?acc=GSE335815.

---

**Supplemental Figure 1:**
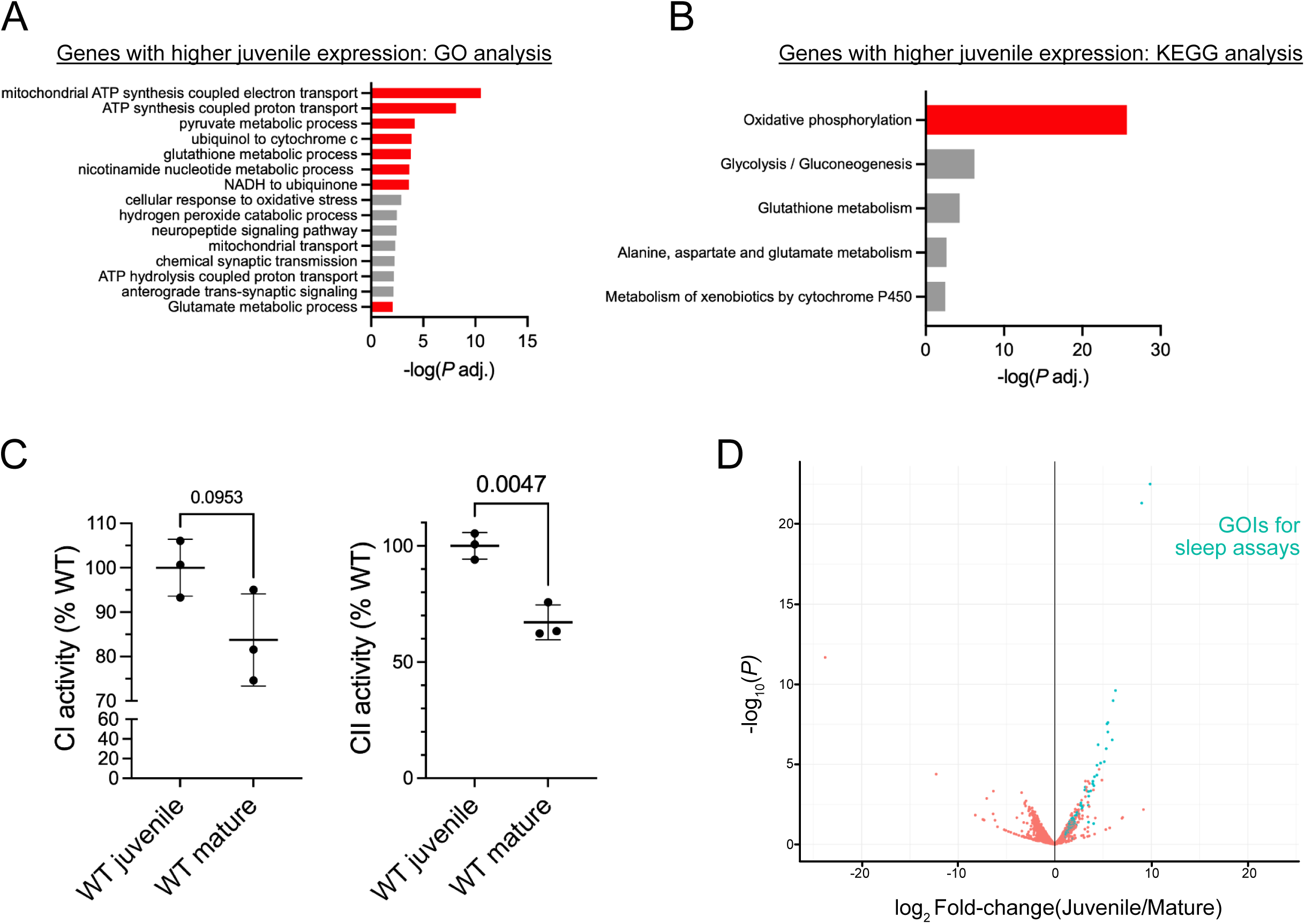
Differential expression analysis of juvenile and mature dopaminergic neurons. (A) GO analysis of differentially expressed genes chosen for RNAi screening. Genes preferentially expressed in juvenile DANs were more likely to encode components of metabolic pathways (examples highlighted in red). (B) KEGG pathway analysis of the same gene set revealed a similar enrichment of metabolic pathways with the highest enrichment mapping to components of the oxidative phosphorylation pathway.)(C) The electron transfer activity of Complexes I and II in mitochondrial fractions isolated from wild-type *iso^31^* juvenile (1 day old) and mature (9-10 day old) adults. Juvenile flies had higher Complex II activity and a trend toward higher Complex I activity. P values computed by Welch’s t-tests. (D) Volcano plot of differential expression showing genes of interest (GOIs, cyan) chosen for the transgenic RNAi screen.

**Supplemental Figure 2:**
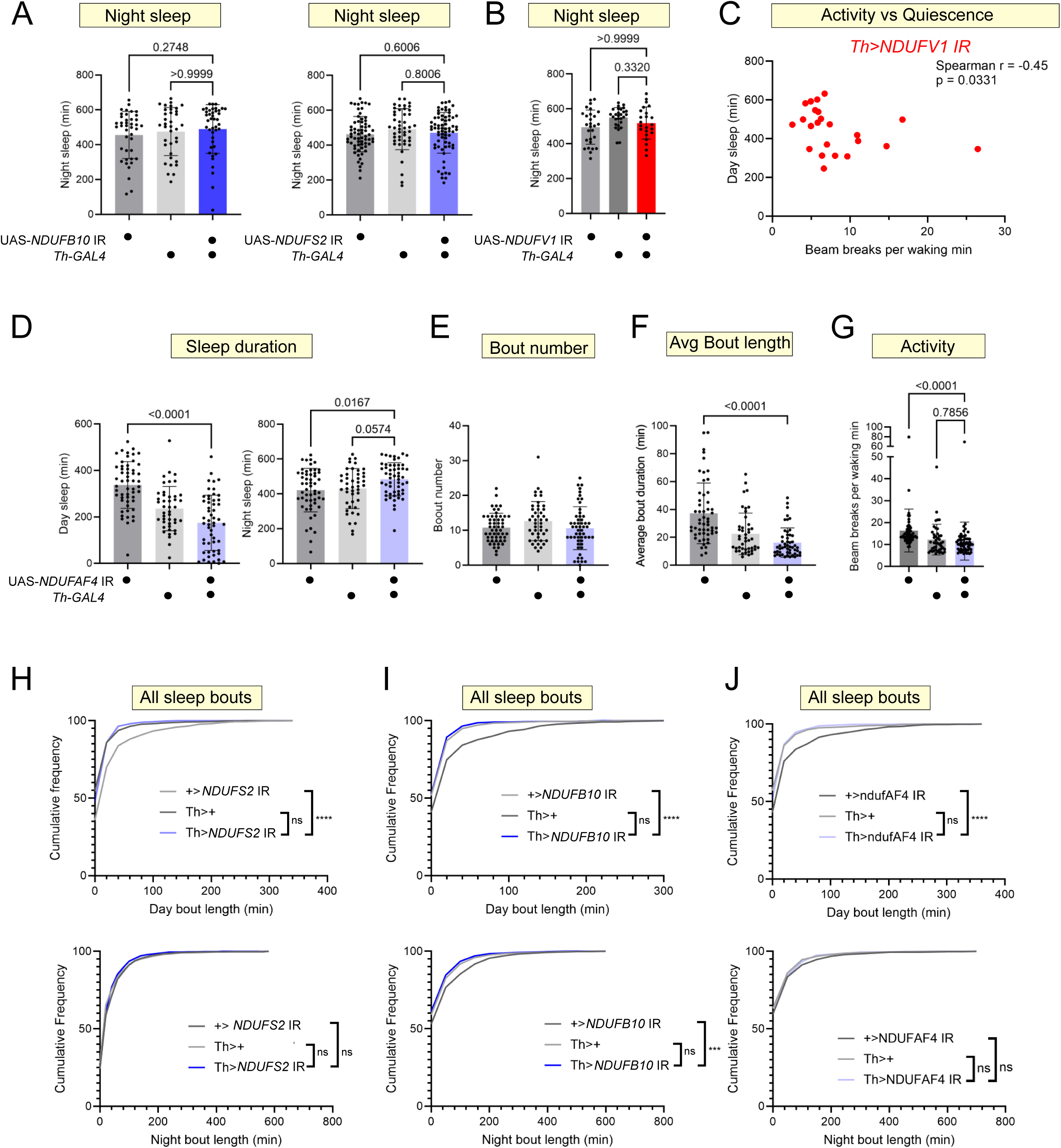
Additional sleep metrics for *Th>*MCI RNAi flies. (A) Night sleep was unaffected in *Th>NDUFB10* IR and *Th>NDUFS2* IR short-sleeping flies. (B) Night sleep was unaffected in *Th>NDUFV1* IR long-sleeping flies. (C) Day sleep duration and waking locomotor activity plotted for each *Th>NDUFV1* IR fly (red dots). The duration of day sleep was significantly correlated with reduced locomotion (Spearman r=-0.45, p=0.0331), indicating variability in sleep can be explained by changes in locomotor behavior. (D-G) Multibeam sleep data for mature adult *Th*>*NDUFAF4* IR flies. Daytime sleep loss was variable in these flies, though many slept far less than controls. (H-J) Cumulative frequency plots of sleep bout duration for short-sleeping Th>MCI RNAi lines. Knock-down adult flies did not show a consistent difference in sleep bout length relative to both control (ns, not significant; *** p<0.001 **** p<0.0001). P values computed using either Kruskal-Wallis tests with Dunn’s multiple-testing comparisons or Welch ANOVA with Dunnett’s T3 multiple-testing comparisons.

**Supplemental Figure 3:**
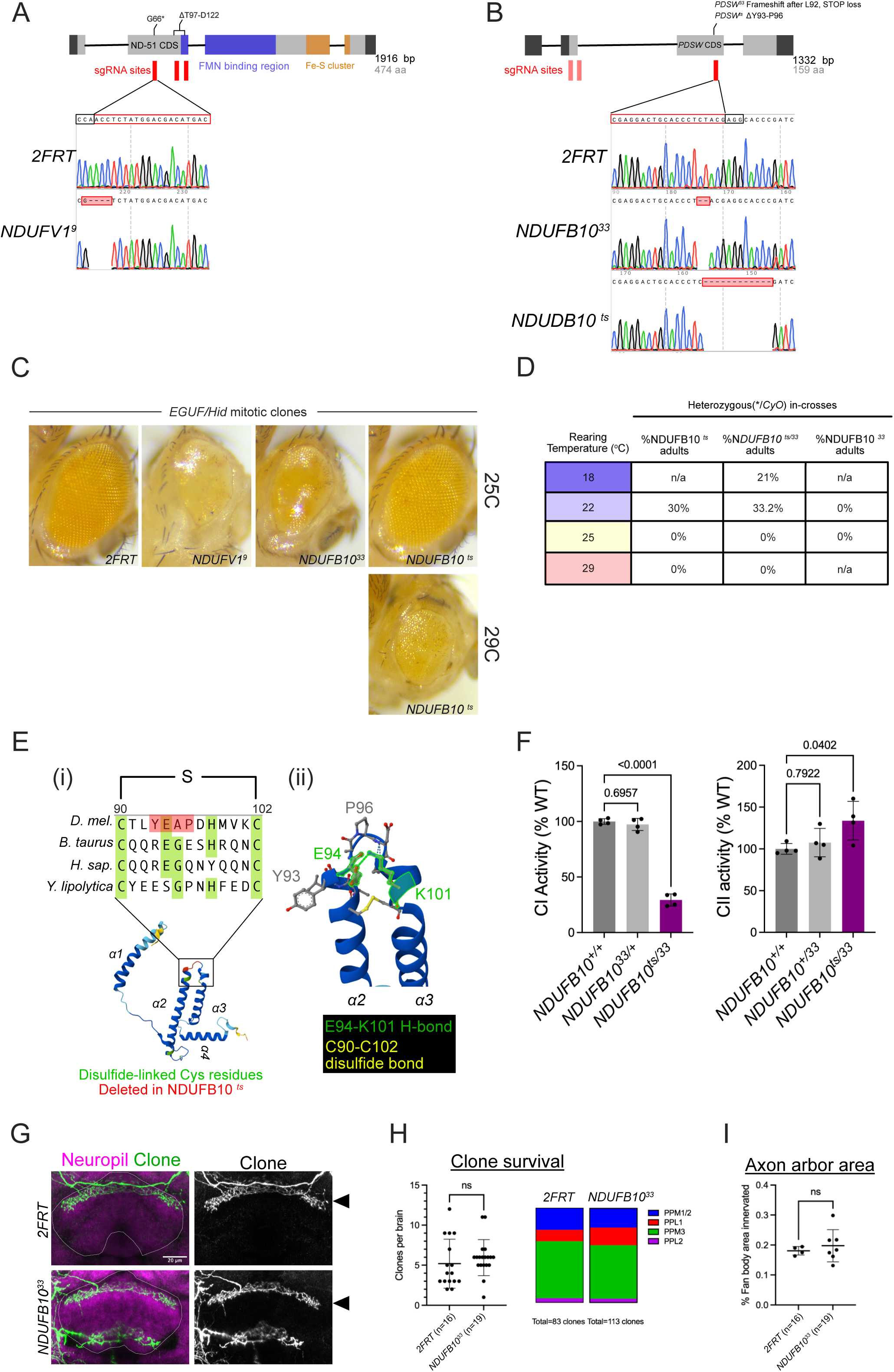
Germline mutants for *NDUFV1* and *NDUFB10* and their phenotypes. (A,B) Position and Sanger chromatograms of germline mutations induced by CRISPR/Cas9. Wild-type sequence derived from the unmutagenized *FRT40A, FRT2A (2FRT*) chromosome. Position of single guide RNAs used shown in red (PAM sequences in black boxes). Shaded red indicates sgRNA that did not induce indels. (C) Mitotic recombination was induced in eye precursor cells. *NDUFB10^33^* (L92fs*?) and *NDUFV1^9^* (G66*, ΔT97-D122) mutants caused small, glossy eyes similar to those reported for other OXPHOS pathway genes. *NDUFB10^ts^* (ΔY93-P96) mutant clones were overtly normal at 25°C but abnormal at 29°C. (D) *NDUFB10^ts^* can rescue the lethality of *NDUFB10^33^* at 22°C and 18°C but not higher temperatures. (E) ClustalOmega alignment of a conserved disulfide bridge in NDUFB10. Both mutant alleles affect a conserved intramolecular disulfide bridge. The NDUFB10*^ts^* allele deletes 4 amino acids predicted by AlphaFold to stabilize the disulfide bonded alpha helices. (F) Spectrophotometric Complex I/II assays in mitochondria isolated from wild-type (*NDUFB10^+/+^*) flies, *NDUFB10^33/+^* heterozygotes, and *NDUFB10^33/ts^* mutants reared at 22°C during development and shifted to 29°C for 1 week to inactivate the NDUFB10^ts^ allele. Mitochondria from the mutants lost ∼80% of wild-type Complex I activity with a compensatory increase in electron flux through Complex II. P values computed using One-way ANOVA’s. (G) Axon terminals of GFP-labeled wild-type (*2FRT*) and *NDUFB10^33^* PPL1 clones projecting to the dorsal fan-shaped body (arrowheads. Neuropil (magenta) labelled by nc82 mAb. (H) The frequency (clone #/brain) and overall distribution of DAN MARCM clones recovered. The *NDUFB10^33^* mutation did not impair clone survival. P values were computed using a Mann-Whitney t-test. (I) The axon arbor area of wild-type and *NDUFB10^33^* clones synapsing onto the dorsal fan-shaped body. The *NDUFB10^33^* mutation did not impair axon pathfinding and innervation of DANs. P value was computed using Welch’s t-test.

**Supplemental Figure 4:**
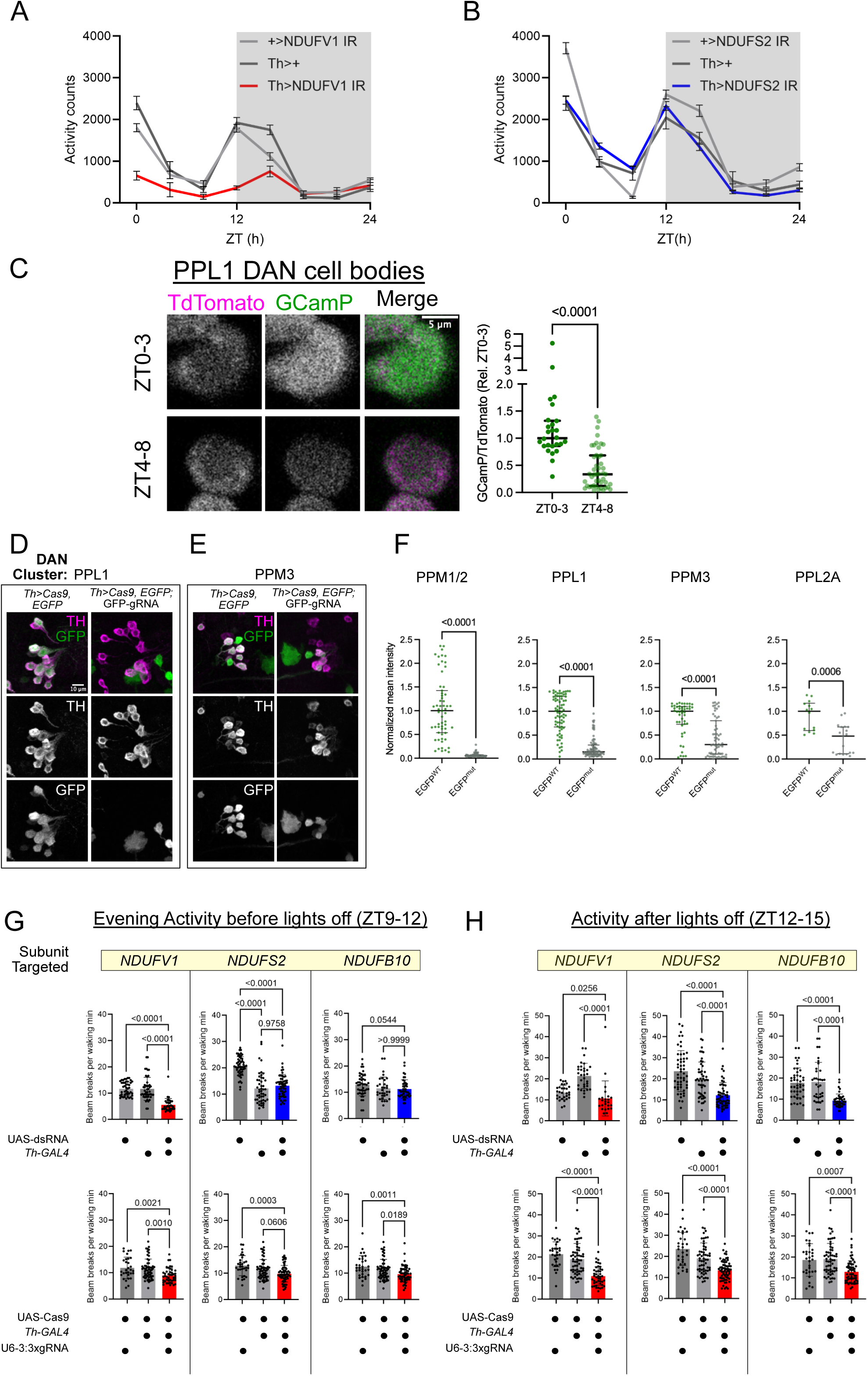
MCI activity in DANs is required for the morning and evening activity peaks. (A,B) Raw activity counts binned into 3h intervals for strong MCI LOF (*Th>NDUFV1* IR, red) and partial MCI LOF (*Th>NDUFB10* IR, blue) flies. Strong MCI, but not partial MCI, LOF severely depressed morning and evening activity peaks. (C) Wild-type PPL1 DANs expressing TdTomato-2A-GCamP6s and imaged *ex vivo*. The steady-state GCamP/TdTomato ratio was higher in the morning (ZT0-3) than the afternoon (ZT4-8) during the afternoon siesta. P value computed by a Mann-Whitney t-test. (D-F) Confirmation of Cas9 mutagenic activity in DANs on the *EGFP* coding sequence. Reduced or loss of EGFP (green) signal indicates Cas9-mediated mutagenesis. Some DAN clusters (PPM1/2 and PPL1) showed highly uniform Cas9 activity, while other clusters (PPM3 and PPL2A) had more heterogeneous activity. Mean EGFP intensity in DAN cell bodies was measured in the absence (green) or presence (grey) of Cas9. Data are normalized to the median of the control. P values computed using Mann-Whitney tests. (G,H) Quantification of evening waking activity for MCI RNAi (top) and MCI somatic knock-out (bottom) flies. Red corresponds to hypoactive conditions, blue to short-sleepers. P values computed using Kruskal-Wallis tests with Dunn’s multiple-testing comparisons and Welch ANOVAs with Dunnett’s T3 multiple-testing comparisons.

**Supplemental Figure 5:**
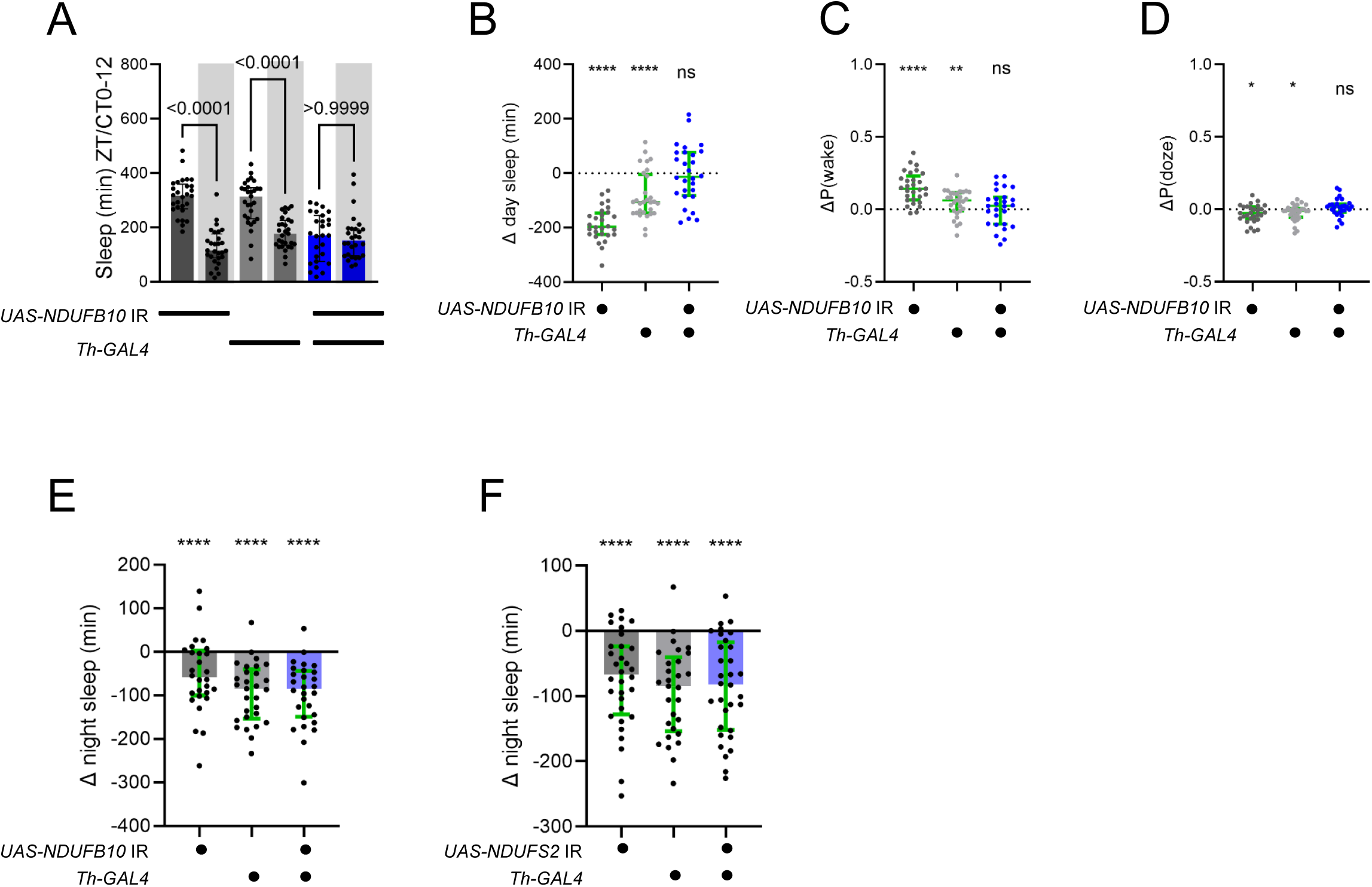
Disinhibition of DA signaling by withdrawal of the light cue modifies wild-type sleep, but not MCI-deficient, flies. A) Sleep during ZT0-12 (lights on) or CT0-12 (no lights on, shaded) for *Th>NDUFB10 IR* mature flies (blue) and controls (grey). Sleep loss in the knock-down flies persisted in constant darkness. Control flies showed a pronounced sleep loss compared to the previous day with lights on. This phenotype was abrogated in *Th>NDUFB10* IR flies. (B) Change in day sleep [Day sleep(DD)-Day sleep (LD)] plotted for each fly in the experiment in B. Values < 0 (dotted line) indicate loss of sleep. *NDUFB10* knock-down abrogated sleep loss due to withdrawal of the light cue. (C)Change in P(wake) [P(wake)(DD)-P(wake)(LD)] plotted for each fly in panel A. Sleep loss upon withdrawal of the light cue was driven by an increase in P(wake)--an effect attenuated *NDUFB10* knock-down. (D) Change in P(doze) [P(doze)(DD)-P(doze)(LD)] plotted for each fly in panel B. Withdrawal of the light cue caused a slight but significant decrease in P(doze) that was not affected by *NDUFS2* knock-down. (E,F) Change in night sleep for *Th>NDUFB10* IR mature flies (blue) and genetic controls (grey) (mean and interquartile range shown). P values calculated by Wilcoxon signed rank tests (* P<0.05, ** P<0.01, **** p<0.0001 compared to Δ=0). Mean and standard deviation plotted for each graph.

**Supplemental Figure 6:**
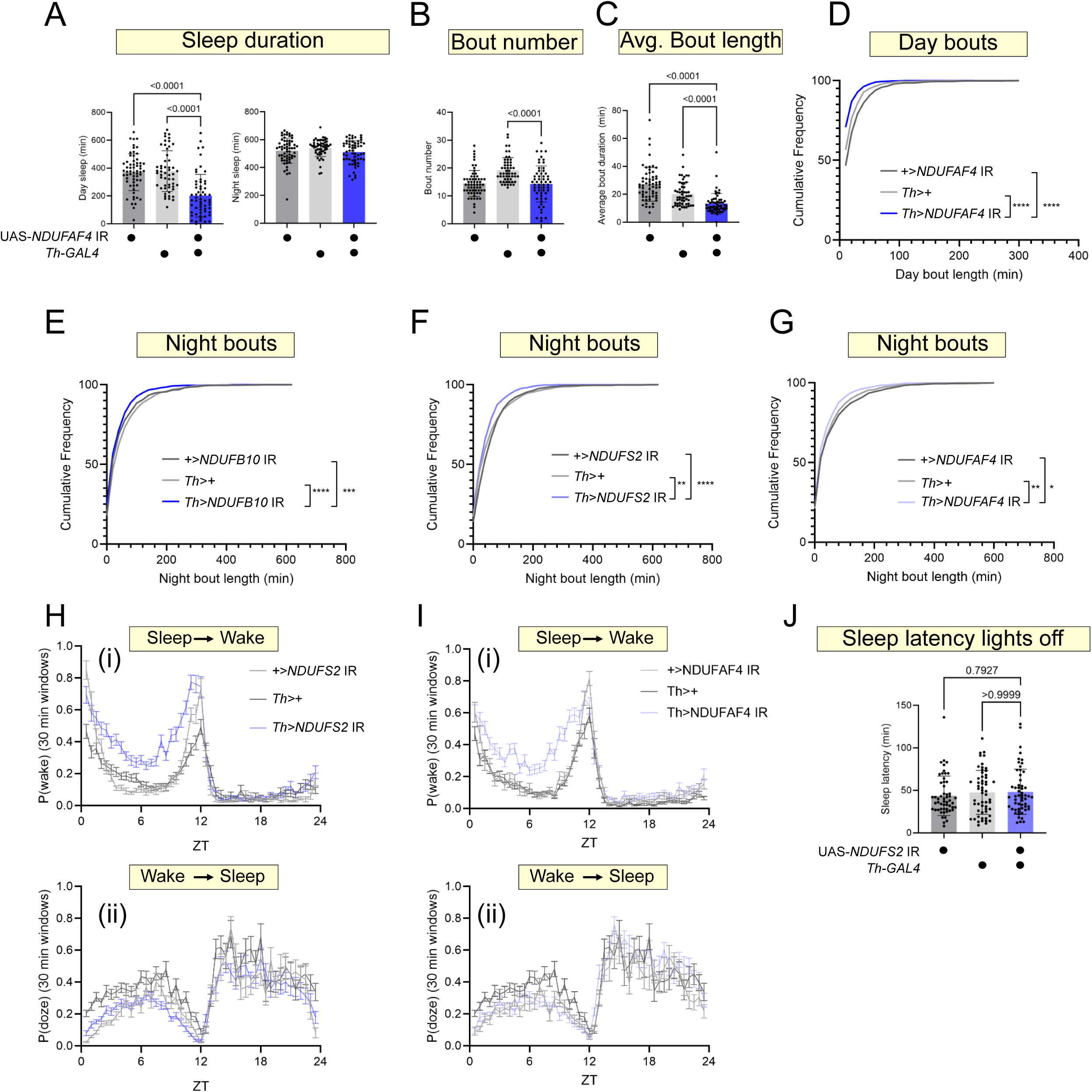
Additional juvenile sleep metrics for partial MCI loss-of-function. (A-D) Juvenile sleep phenotypes for *Th>NDUFAF4* IR juvenile flies and controls. Knock-down flies exhibited significant daytime sleep loss (A) driven by reduced bout duration (C, D, **** p<0.0001 Kruskal-Wallis with Dunn’s multiple comparisons test). (E-G) Cumulative frequency plots for durations of nighttime sleep bouts of partial MCI knock-down flies. Juvenile flies for all three knock-down conditions showed significantly reduced bout length during the night (** p<0.01, *** p<0.001, **** p<0.0001 Kruskal-Wallis with Dunn’s multiple testing comparisons). (H) P(wake) (i) and P(doze) (ii) measurements (average and SEM) for 30 min windows across ZT0-24 for juvenile *Th>NDUFS2* IR juvenile flies (blue) and controls (grey). Knock-down flies showed significantly elevated P(wake) across ZT0-12, indicating sleep loss during this period was driven by reduced sleep depth. P(doze) was only visibly reduced compared to both controls during ZT6-12. (I) P(wake) (i) and P(doze) (ii) measurements (average and SEM) for 30 min windows across ZT0-24 for juvenile *Th>NDUFAF4* IR juvenile flies (blue) and controls (grey). Juvenile *Th>NDUFAF4* IR sleep loss was driven by elevated P(wake). (J) Time to first sleep bout after lights-off (ZT12) for juvenile *Th>NDUFS2* IR juvenile flies and genetic controls, showing no difference across genotypes. (K) Waking locomotor activity during morning (ZT0-3) for *Th>NDUFS2* IR juvenile flies and genetic controls. Despite increased sleep latency during this period (see Fig 4), knock-down flies did not exhibit elevated locomotor activity, indicating prolonged sleep latency is not a secondary consequence of hyperactivity. P values computed using either Kruskal-Wallis tests with Dunn’s multiple-testing comparisons or Welch ANOVA with Dunnett’s T3 multiple-testing comparisons.

**Supplemental Figure 7:**
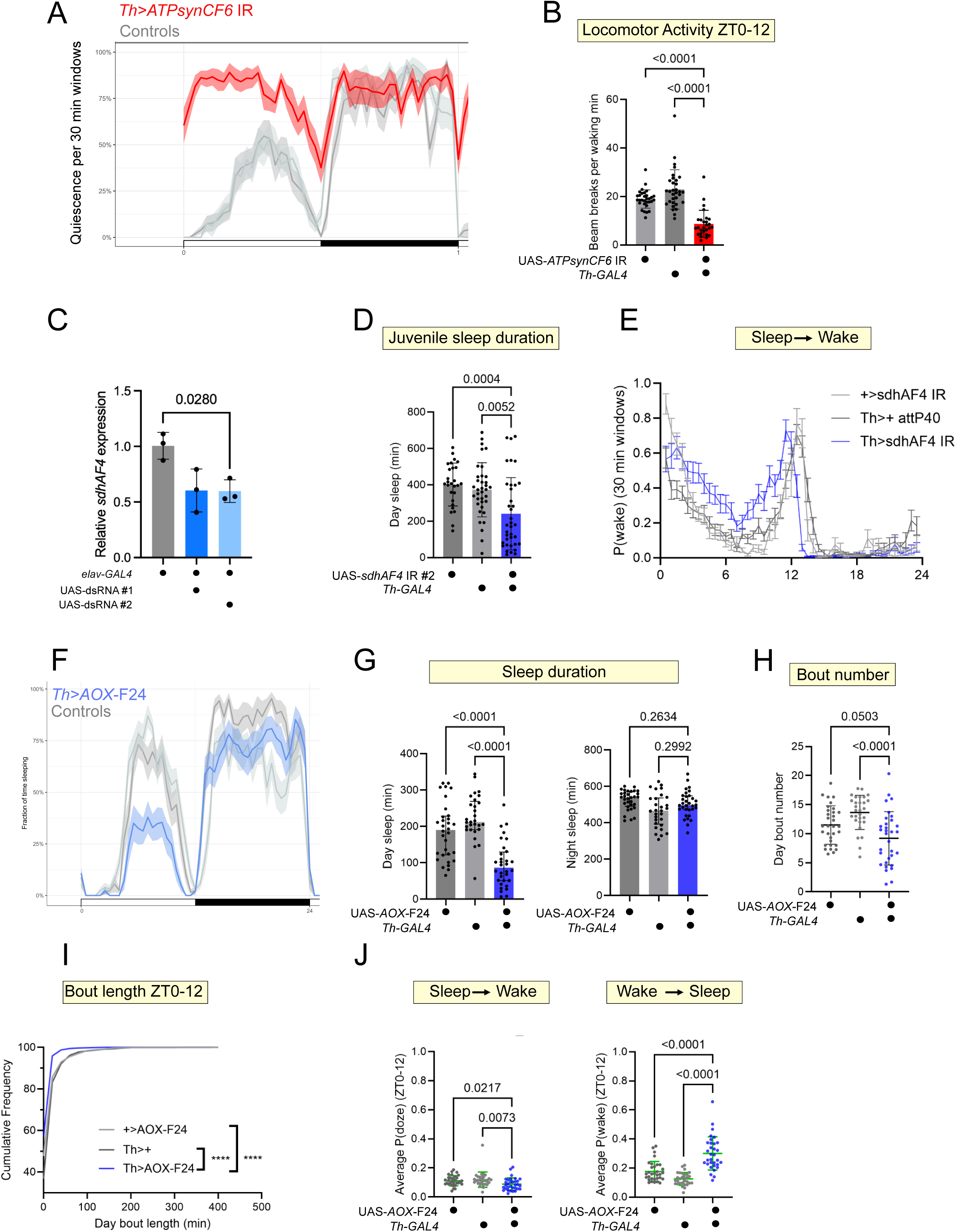
Depletion of dopaminergic CoQH2 by multiple, independent means caused sleep loss. (A) Quiescence per 30 min windows (mean with 95% confidence interval shaded) for *Th>ATPsynCF6* IR mature adult flies (red), expressing a dsRNA against an essential structural component of the F(1)-F(0) ATPase, and genetic controls (grey). The dramatic increase in daytime quiescence reflects severely reduced locomotion (B, p<0.0001 Welch’s ANOVA with Dunnet’s T3 multiple comparisons test). (C) Quantitative PCR of whole-head cDNA from *elav>sdhAF4* IR flies (blue) and the landing site control (grey). The two independent short-sleeping RNAi lines targeting *sdhAF4* comparably reduced *sdhAF4* mRNA levels. P value computed using Welch ANOVA and Dunnett’s T3 multiple comparisons test. (D) Juvenile sleep duration during ZT0-12 for *Th>sdhAF4* IR #2 (blue) and controls (grey) assayed using the multibeam activity monitoring system, confirming sleep loss phenotype reported in primary screen. (E) P(wake) plotted in 30 min windows (average with SEM) for juvenile *Th>sdhAF4* IR #2 (blue) and controls (grey) showing a persistently elevated P(wake) during ZT0-12, indicative of increased transitions from inactive to active—a phenotype is identical to partial MCI inhibition. (F) Sleep per 30 min windows (meanwith 95% confidence interval shaded) of *Th>AOX-F24* mature adult flies (blue) expressing an independent P element insertion of UAS-*AOX* and genetic controls (grey). (G) Day and night sleep duration for mature adult *Th>AOX-F24* flies (blue). Compared to the insertion used in Fig 5, UAS-*AOX-F24* flies showed comparable sleep loss during ZT0-12 but no effect on sleep during ZT12-24. P values computed using a Kruskal-Wallis test with Dunn’s multiple testing comparisons. (H, I) Sleep bout metrics for *Th>AOX-F24.* Sleep loss was driven by both reduced bout number and bout length (**** p<0.0001, Kruskal-Wallis test with Dunn’s multiple testing comparisons). (J, K) P(doze) and P(wake) averaged across ZT0-12 for *Th>AOX-F24* flies (blue) and genetic controls (grey). As with UAS-AOX transgene used in Fig 5, *Th>AOX-F24* flies showed a substantial increase in P(wake), and a modest decrease in P(doze), during ZT0-12. P values computed using Kruskal-Wallis tests with Dunn’s multiple-testing comparisons.

## Literature Cited

1. Kayser MS, Biron D. Sleep and Development in Genetically Tractable Model Organisms. Genetics. 2016;203(1):21–33.

2. Roffwarg HP, Muzio JN, Dement WC. Ontogenetic Development of the Human Sleep-Dream Cycle. Science. 1966;152(3722):604–19.

3. Dashti HS, Jones SE, Wood AR, Lane JM, van Hees VT, Wang H, et al. Genome-wide association study identifies genetic loci for self-reported habitual sleep duration supported by accelerometer-derived estimates. Nature Communications. 2019;10(1):1100.

4. Bian WJ, Brewer CL, Kauer JA, de Lecea L. Adolescent sleep shapes social novelty preference in mice. Nat Neurosci. 2022;25(7):912–23.

5. Kayser MS, Yue Z, Sehgal A. A critical period of sleep for development of courtship circuitry and behavior in Drosophila. Science. 2014;344(6181):269–74.

6. Gay SM, Chartampila E, Lord JS, Grizzard S, Maisashvili T, Ye M, et al. Developing forebrain synapses are uniquely vulnerable to sleep loss. Proc Natl Acad Sci U S A. 2024;121(44):e2407533121.

7. Jones CE, Opel RA, Kaiser ME, Chau AQ, Quintana JR, Nipper MA, et al. Early-life sleep disruption increases parvalbumin in primary somatosensory cortex and impairs social bonding in prairie voles. Sci Adv. 2019;5(1):eaav5188.

8. Veatch OJ, Maxwell-Horn AC, Malow BA. Sleep in Autism Spectrum Disorders. Curr Sleep Med Rep. 2015;1(2):131–40.

9. Veatch OJ, Sutcliffe JS, Warren ZE, Keenan BT, Potter MH, Malow BA. Shorter sleep duration is associated with social impairment and comorbidities in ASD. Autism Res. 2017;10(7):1221–38.

10. Dilley LC, Vigderman A, Williams CE, Kayser MS. Behavioral and genetic features of sleep ontogeny in Drosophila. Sleep. 2018;41(7).

11. Shaw PJ, Cirelli C, Greenspan RJ, Tononi G. Correlates of sleep and waking in Drosophila melanogaster. Science. 2000;287(5459):1834–7.

12. Chakravarti Dilley L, Szuperak M, Gong NN, Williams CE, Saldana RL, Garbe DS, et al. Identification of a molecular basis for the juvenile sleep state. Elife. 2020;9.

13. Agip A-NA, Chung I, Sanchez-Martinez A, Whitworth AJ, Hirst J. Cryo-EM structures of mitochondrial respiratory complex I from Drosophila melanogaster. eLife. 2023;12:e84424.

14. Murari A, Rhooms SK, Garcia C, Liu T, Li H, Mishra B, et al. Dissecting the concordant and disparate roles of NDUFAF3 and NDUFAF4 in mitochondrial complex I biogenesis. iScience. 2021;24(8):102869.

15. Arroum T, Borowski MT, Marx N, Schmelter F, Scholz M, Psathaki OE, et al. Loss of respiratory complex I subunit NDUFB10 affects complex I assembly and supercomplex formation. Biol Chem. 2023;404(5):399–415.

16. Stroud DA, Surgenor EE, Formosa LE, Reljic B, Frazier AE, Dibley MG, et al. Accessory subunits are integral for assembly and function of human mitochondrial complex I. Nature. 2016;538(7623):123–6.

17. Port F, Chen H-M, Lee T, Bullock SL. Optimized CRISPR/Cas tools for efficient germline and somatic genome engineering in Drosophila. Proceedings of the National Academy of Sciences. 2014;111(29):E2967–E76.

18. Rimal S, Tantray I, Li Y, Pal Khaket T, Li Y, Bhurtel S, et al. Reverse electron transfer is activated during aging and contributes to aging and age-related disease. EMBO Reports. 2023;24(4):EMBR202255548.

19. Liu Q, Liu S, Kodama L, Driscoll MR, Wu MN. Two dopaminergic neurons signal to the dorsal fan-shaped body to promote wakefulness in Drosophila. Curr Biol. 2012;22(22):2114–23.

20. Ueno T, Tomita J, Tanimoto H, Endo K, Ito K, Kume S, et al. Identification of a dopamine pathway that regulates sleep and arousal in Drosophila. Nat Neurosci. 2012;15(11):1516–23.

21. Liao TS, Call GB, Guptan P, Cespedes A, Marshall J, Yackle K, et al. An efficient genetic screen in Drosophila to identify nuclear-encoded genes with mitochondrial function. Genetics. 2006;174(1):525–33.

22. Mandal S, Guptan P, Owusu-Ansah E, Banerjee U. Mitochondrial regulation of cell cycle progression during development as revealed by the tenured mutation in Drosophila. Dev Cell. 2005;9(6):843–54.

23. Stowers RS, Schwarz TL. A genetic method for generating Drosophila eyes composed exclusively of mitotic clones of a single genotype. Genetics. 1999;152(4):1631– 9.

24. Friederich MW, Erdogan AJ, Coughlin CR, 2nd, Elos MT, Jiang H, O’Rourke CP, et al. Mutations in the accessory subunit NDUFB10 result in isolated complex I deficiency and illustrate the critical role of intermembrane space import for complex I holoenzyme assembly. Hum Mol Genet. 2017;26(4):702–16.

25. Jumper J, Evans R, Pritzel A, Green T, Figurnov M, Ronneberger O, et al. Highly accurate protein structure prediction with AlphaFold. Nature. 2021;596(7873):583–9.

26. Mitchell DV, Iadarola DM, Mathew ND, Keith K, Seiler C, Yu S, et al. ndufs2(-/-) zebrafish have impaired survival, neuromuscular activity, morphology, and one-carbon metabolism treatable with folic acid. bioRxiv. 2025.

27. Lee T, Luo L. Mosaic analysis with a repressible cell marker (MARCM) for Drosophila neural development. Trends Neurosci. 2001;24(5):251–4.

28. Aimon S, Katsuki T, Jia T, Grosenick L, Broxton M, Deisseroth K, et al. Fast near-whole–brain imaging in adult Drosophila during responses to stimuli and behavior. PLOS Biology. 2019;17(2):e2006732.

29. Le JQ, Ma D, Dai X, Rosbash M. Light and dopamine impact two circadian neurons to promote morning wakefulness. Curr Biol. 2024;34(17):3941–54.e4.

30. Chen X, Perry S, Fan Z, Wang B, Loxterkamp E, Wang S, et al. Tissue-specific knockout in the Drosophila neuromuscular system reveals ESCRT’s role in formation of synapse-derived extracellular vesicles. PLOS Genetics. 2024;20(10):e1011438.

31. Shang Y, Haynes P, Pírez N, Harrington KI, Guo F, Pollack J, et al. Imaging analysis of clock neurons reveals light buffers the wake-promoting effect of dopamine. Nature Neuroscience. 2011;14(7):889–95.

32. Van Vranken JG, Bricker DK, Dephoure N, Gygi SP, Cox JE, Thummel CS, et al. SDHAF4 promotes mitochondrial succinate dehydrogenase activity and prevents neurodegeneration. Cell Metab. 2014;20(2):241–52.

33. Fernandez-Ayala DJM, Sanz A, Vartiainen S, Kemppainen KK, Babusiak M, Mustalahti E, et al. Expression of the Ciona intestinalis Alternative Oxidase (AOX) in Drosophila Complements Defects in Mitochondrial Oxidative Phosphorylation. Cell Metabolism. 2009;9(5):449–60.

34. Osterwalder T, Yoon KS, White BH, Keshishian H. A conditional tissue-specific transgene expression system using inducible GAL4. Proceedings of the National Academy of Sciences. 2001;98(22):12596–601.

35. Kume K, Kume S, Park SK, Hirsh J, Jackson FR. Dopamine is a regulator of arousal in the fruit fly. J Neurosci. 2005;25(32):7377–84.

36. Tryphena KP, Nikhil US, Pinjala P, Srivastava S, Singh SB, Khatri DK. Mitochondrial Complex I as a Pathologic and Therapeutic Target for Parkinson’s Disease. ACS Chemical Neuroscience. 2023;14(8):1356–68.

37. Musiek ES, Holtzman DM. Mechanisms linking circadian clocks, sleep, and neurodegeneration. Science. 2016;354(6315):1004–8.

38. Summa KC, Jiang P, González-Rodríguez P, Huang X, Lin X, Vitaterna MH, et al. Disrupted sleep-wake regulation in the MCI-Park mouse model of Parkinson’s disease. npj Parkinson’s Disease. 2024;10(1):54.

39. Oswald MCW, Brooks PS, Zwart MF, Mukherjee A, West RJH, Giachello CNG, et al. Reactive oxygen species regulate activity-dependent neuronal plasticity in Drosophila. eLife. 2018;7:e39393.

40. Li Z, Ji G, Neugebauer V. Mitochondrial reactive oxygen species are activated by mGluR5 through IP3 and activate ERK and PKA to increase excitability of amygdala neurons and pain behavior. J Neurosci. 2011;31(3):1114–27.

41. Beckhauser TF, Francis-Oliveira J, De Pasquale R. Reactive Oxygen Species: Physiological and Physiopathological Effects on Synaptic Plasticity. J Exp Neurosci. 2016;10(Suppl 1):23–48.

42. Tamsett TJ, Picchione KE, Bhattacharjee A. NAD+ activates KNa channels in dorsal root ganglion neurons. J Neurosci. 2009;29(16):5127–34.

43. Rangaraju V, Calloway N, Ryan Timothy A. Activity-Driven Local ATP Synthesis Is Required for Synaptic Function. Cell. 2014;156(4):825–35.

44. Sarnataro R, Velasco CD, Monaco N, Kempf A, Miesenböck G. Mitochondrial origins of the pressure to sleep. Nature. 2025;645(8081):722–8.

45. Haynes PR, Pyfrom ES, Li Y, Stein C, Cuddapah VA, Jacobs JA, et al. A neuron-glia lipid metabolic cycle couples daily sleep to mitochondrial homeostasis. Nat Neurosci. 2024;27(4):666–78.

46. Kempf A, Song SM, Talbot CB, Miesenböck G. A potassium channel β-subunit couples mitochondrial electron transport to sleep. Nature. 2019;568(7751):230–4.

47. Ni JQ, Liu LP, Binari R, Hardy R, Shim HS, Cavallaro A, et al. A Drosophila resource of transgenic RNAi lines for neurogenetics. Genetics. 2009;182(4):1089–100.

48. Perkins LA, Holderbaum L, Tao R, Hu Y, Sopko R, McCall K, et al. The Transgenic RNAi Project at Harvard Medical School: Resources and Validation. Genetics. 2015;201(3):843–52.

49. Geissmann Q, Garcia Rodriguez L, Beckwith EJ, Gilestro GF. Rethomics: An R framework to analyse high-throughput behavioural data. PLOS ONE. 2019;14(1):e0209331.

50. Vecsey CG, Koochagian C, Porter MT, Roman G, Sitaraman D. Analysis of Sleep and Circadian Rhythms from Drosophila Activity-Monitoring Data Using SCAMP. Cold Spring Harb Protoc. 2024;2024(11):pdb.prot108182.

51. Tricoire H, Battisti V, Trannoy S, Lasbleiz C, Pret AM, Monnier V. The steroid hormone receptor EcR finely modulates Drosophila lifespan during adulthood in a sex-specific manner. Mech Ageing Dev. 2009;130(8):547–52.

52. Ye J, Coulouris G, Zaretskaya I, Cutcutache I, Rozen S, Madden TL. Primer-BLAST: a tool to design target-specific primers for polymerase chain reaction. BMC Bioinformatics. 2012;13:134.

53. Groen CM, Windebank AJ. Isolation of Intact Mitochondria From Drosophila melanogaster and Assessment of Mitochondrial Respiratory Capacity Using Seahorse Analyzer. Bio Protoc. 2025;15(3):e5180.

54. Graeff R, Lee HC. A novel cycling assay for cellular cADP-ribose with nanomolar sensitivity. Biochemical Journal. 2002;361(2):379–84.

55. Mukherjee S, Velázquez Aponte RA, Perry CE, Lee WD, Janssen KA, Niere M, et al. Hepatocyte mitochondrial NAD+ content is limiting for liver regeneration. Nature Metabolism. 2025;7(12):2424–37.

